# The function of CozE proteins is linked to lipoteichoic acid biosynthesis in *Staphylococcus aureus*

**DOI:** 10.1101/2023.10.20.563254

**Authors:** Maria Disen Barbuti, Elisabeth Lambert, Ine Storaker Myrbråten, Adrien Ducret, Gro Anita Stamsås, Linus Wilhelm, Xue Liu, Zhian Salehian, Jan-Willem Veening, Daniel Straume, Christophe Grangeasse, Camilo Perez, Morten Kjos

**Affiliations:** Faculty of Chemistry, Biotechnology and Food Science, Norwegian University of Life Sciences, Ås, Norway; Biozentrum, University of Basel, Basel, Switzerland; Molecular Microbiology and Structural Biochemistry, CNRS UM 5086, Université de Lyon, Lyon, France; Department of Pathogen, Biology, International Cancer Center, Shenzhen University Medical School, Shenzhen, Guangdong, 518055, China; Department of Fundamental Microbiology, University of Lausanne, Switzerland

## Abstract

To maintain cell integrity and facilitate cell division in *Staphylococcus aureus*, a well-coordinated interplay between membrane biogenesis, peptidoglycan formation, and teichoic acid synthesis is crucial. However, the molecular mechanisms and regulatory pathways that underpin their coordination are still poorly understood. CozE constitute a conserved family of membrane proteins implicated in cell division via regulation of penicillin binding proteins. It has been shown that the two staphylococcal *cozE* genes (*cozEa* and *cozEb*) constitute a synthetic lethal gene pair. Depletion of CozEa and CozEb simultaneously in *S. aureus* resulted in severely defective cell division phenotypes, reminiscent of cell lacking lipoteichoic acid (LTA). Indeed, we demonstrate that there is an intricate interplay between CozE, biosynthesis of LTA, and membrane homeostasis in *S. aureus*. By screening for potential genetic links, we establish that there is synthetic lethal relationship between CozE and UgtP, the enzyme synthesizing the LTA glycolipid anchor Glc_2_DAG. On the contrary, in cells lacking LtaA, the flippase of Glc_2_DAG, the essentiality of CozEa and CozEb was alleviated. Furthermore, by immunoblotting, we found that CozEb plays a unique role in controlling LTA polymer length and stability. Using reconstituted proteoliposomes, we also demonstrated that CozE proteins modulate the glycolipid flipping activity of LtaA *in vitro*. Together, the results demonstrate a new function of CozE proteins, facilitating proper membrane homeostasis and LTA biosynthesis in *S. aureus*.

## Introduction

*Staphylococcus aureus* is a Gram-positive, opportunistic pathogen which is responsible for a wide range of infectious diseases in humans and animals, including skin and soft tissue infections, bloodstream infections, and infections associated with medical implant devices. This is made possible by the plethora of virulence factors produced by *S. aureus*. Among the major factors contributing to staphylococcal colonization, infection, and immune evasion are the anionic teichoic acid (TA) polymers (1, 2). Together with peptidoglycan, TAs are the main constituents of the staphylococcal cell wall. Interestingly, TAs do not only influence the susceptibility of *S. aureus* to antibiotics, but they also regulate it (1, 3), making the TA biosynthetic pathways attractive as potential anti-virulence and antibiotic targets.

Staphylococcal TAs are mainly composed of repeating units of ribitol phosphate (RboP) or glycerol phosphate (GroP), that are either covalently linked to peptidoglycan (wall teichoic acids, WTAs) or anchored to the cytoplasmic membrane (lipoteichoic acids, LTAs). Although LTA is more important than WTA for cell viability, it is possible to create deletion mutants of genes involved in either pathway. However, it is not possible to delete both pathways simultaneously (4, 5). Staphylococcal LTAs consist of poly-GroP chains that are associated with the cytoplasmic membrane through the glycolipid anchor diglucosyl-diacylglycerol (Glc_2_DAG) (**Fig. 3A**). Glc_2_DAG is synthesized in the cytoplasm by the glycosyltransferase UgtP (also called YpfP), which transfers two glucose moieties from uridine diphosphate glucose (UDP-Glc) to DAG (6). Glc_2_DAG is translocated to the outer membrane leaflet by the multi-membrane spanning protein LtaA (7, 8). Lastly, the LTA synthase, LtaS, polymerizes the poly-GroP backbone chain by transferring GroP units, derived from the head group of phosphatidylglycerol (PG), to the Glc_2_DAG, on the outside surface of the membrane (9), leaving extracellular DAG as a by-product. LTA polymers are often further modified by D-alanylation, carried out by the DltABCD system, and/or glycosylation, which modulates their properties and functions (10, 11).

The LTA synthase LtaS is considered essential for growth in *S. aureus*, as deletion of *ltaS*, in which LTA synthesis is completely abolished, is associated with suppressor mutations (12, 13). Cells with deletion of the genes required for synthesis and flipping of the glycolipid anchor (Δ*ugtP* and Δ*ltaA*, respectively), are however still viable (6, 14). In Δ*ugtP* cells, which completely lack the Glc_2_DAG anchor, LtaS can initiate LTA synthesis directly on PG (6, 15). Δ*ltaA* mutants have been demonstrated to produce a mixture of LTAs linked to both PG and Glc_2_DAG (7); hence, there must exist an unidentified mechanism that can translocate Glc_2_DAG, produced by UgtP, to the outer membrane leaflet in the absence of LtaA. In Δ*ugtP* or Δ*ltaA* cells, LTA length control is lost, resulting in cells that produce PG-linked LTA polymers which are abnormally long and less stable (7, 16, 17).

Importantly, several studies suggest a tight link between LTA synthesis and cell division in *S. aureus*. Deletion of genes in the LTA biosynthetic pathway results in enlarged cells with severe division and morphological defects, suggested to be caused by changes in LTA length and abundance (6, 14, 15, 18). Furthermore, UgtP, LtaA, and LtaS, have all been shown to interact with each other, as well as multiple cell division and cell wall synthesis proteins (e.g., EzrA, FtsA, FtsW, and PBP1-PBP4) (19). LtaS has also been demonstrated to mainly accumulate at the septum in *S. aureus*, indicating that LTA synthesis predominantly occurs at the division site (19). Together this suggest that LTA biosynthesis is tightly coordinated with peptidoglycan synthesis and other processes during the staphylococcal cell cycle (20).

CozE (<u>c</u>oordinator <u>o</u>f <u>z</u>onal <u>e</u>longation) belongs to a family of multi-transmembrane proteins that are broadly distributed across the bacterial kingdom (21). CozE (also referred to as CozEa) was first studied in *Streptococcus pneumoniae*, where it was shown to direct localization of peptidoglycan synthesis possibly via interactions with the bifunctional class A PBP1a and the MreCD complex involved in cell morphogenesis (21–23). Later studies have also indicated functional interactions with additional morphogenesis factors, such as RodZ (24). A CozE paralog in *S. pneumoniae*, named CozEb, has also been found to be part of the same complex as CozE (24, 25), and there seems to be a complex interplay between the two paralogs (24, 25); individual deletions of *cozE* or *cozEb* in *S. pneumoniae* generated different phenotypes with regard to cell shape and growth inhibition, and while CozEb was not required for correct localization of PBP1a, overexpression of this protein could compensate for deletion of *cozE*, suppressing both growth and morphology defects.

In *S. aureu*s, the two CozE-paralogs, CozEa and CozEb, have been studied in the strain SH1000, where they were found to be important for proper cell division (26). While neither *cozEa* nor *cozEb* was essential when deleted individually, a synthetic lethal phenotype was observed; a double deletion strain could not be obtained and knockdown of *cozEa* in a Δ*cozEb* background (or *vice versa*) resulted in significantly reduced growth, aberrant septal placement, distorted cell morphologies, frequent cell lysis, and a non-homogeneous nucleoid staining (26). CozEa and CozEb were found to interact with, and modulate the localization of the cell division regulator EzrA, suggesting that this interaction may be important for the coordination of cell division in this bacterium (26).

In this work, we demonstrate that there is a functional link between biosynthesis of LTA and CozE proteins in *S. aureus*. We show that the two CozE proteins have unique functionalities, as CozEb, but not CozEa, modulates the length and stability of LTAs and the flipping activity of LtaA *in vitro*. The results presented here give insights into hitherto unknown functions of the broadly distributed CozE proteins.

## Results

### CozEa and CozEb affect cell division across different *Staphylococcus aureus* strains

In a previous work, we showed that *cozEa* and *cozEb* in *S. aureus* SH1000 were synthetic lethal and possessed overlapping effects on growth and cell morphology (26). To investigate the functional conservation of *cozEa* and *cozEb* across different *S. aureus* strains and to characterize the phenotypes in more detail, Δ*cozEa::spc* and Δ*cozEb::spc* mutants were constructed in the methicillin-sensitive *S. aureus* (MSSA) strain NCTC8325-4, while the *cozEa::Tn* and *cozEb::Tn* mutants in the community-associated methicillin-resistant *S. aureus* (CA-MRSA) USA300 strain JE2 were obtained from the Nebraska collection (27). Neither of the single deletions resulted in any growth defects under the conditions tested, and no obvious morphological defects were found by microscopy analysis (**Fig. 1A-B**, **Fig. S1A**, **Fig. S2A-B**). The cell sizes were not severely altered, although the JE2 Δ*cozEa* cells, as well as both NCTC8325-4 *cozE* mutants, on average were slightly smaller than their wild-types (**Fig. 1B-C**, **Fig. S2B-C**). Likewise, cell cycle phase distribution analysis (performed on cells with fluorescent vancomycin labelled cell wall; **Fig. 1B**, **Fig S2B**) (28) did not reveal differences between mutant and wild-type cells (**Fig. 1D-E**, **Fig. S2D**). Thus, the *cozEa* and *cozEb* mutants in JE2 and NCTC8325-4 were similar to their respective wild-types, which is consistent with the results from *S. aureus* SH1000 (26).

**Fig. 1.**
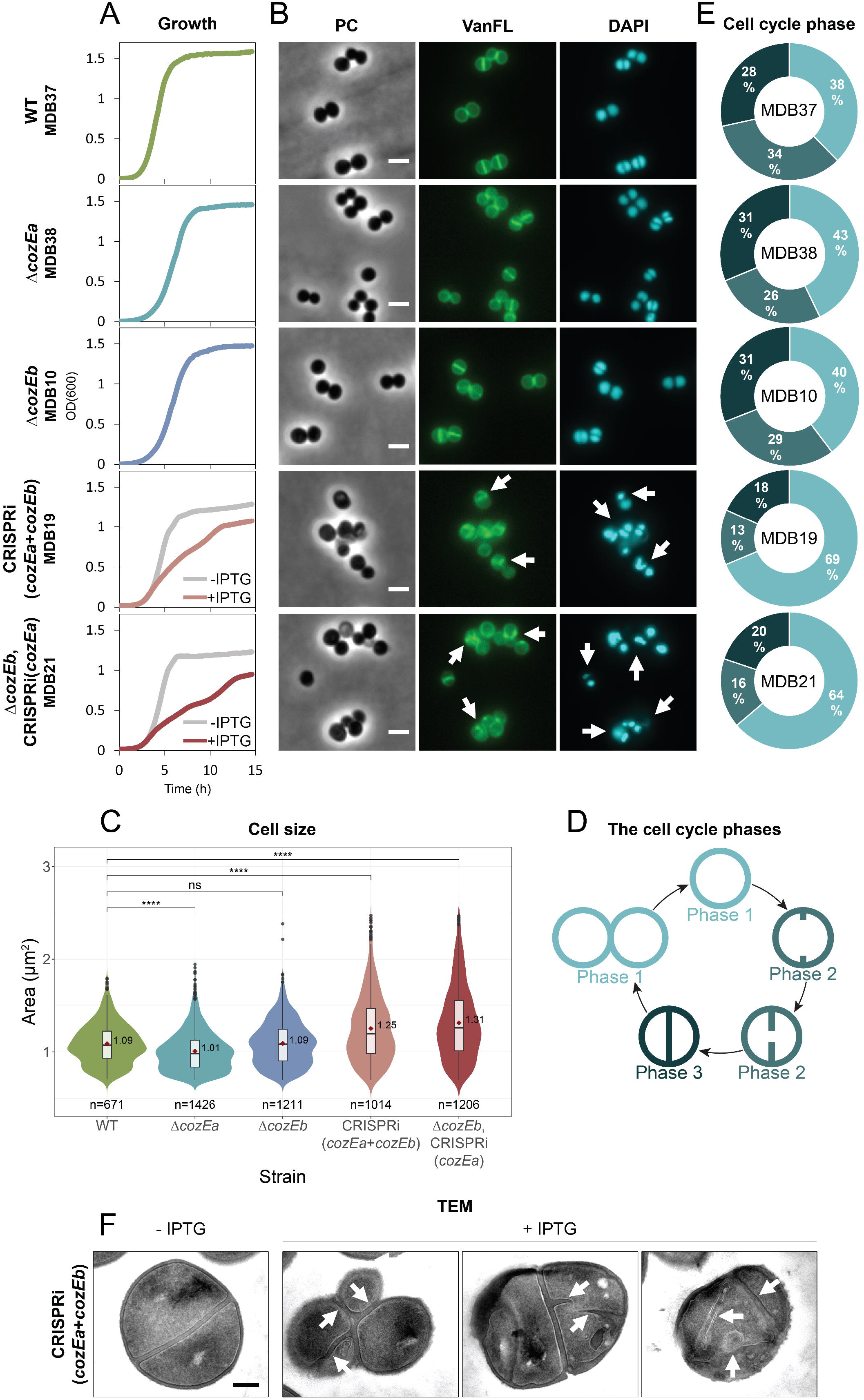
Morphological and cell cycle analysis of single and double *cozE* mutants in *S. aureus* JE2. (**A**) Growth curves of JE2 wild-type (MDB37), Δ*cozEa*::Tn (MDB38), and Δ*cozEb*::Tn (MDB10), as well as of a CRISPRi double knockdown strain (CRISPRi(*cozEa+cozEb*), MDB19) and a combined knockout/knockdown strain (Δ*cozEb*+CRISPRi(*cozEa*), MDB21) in BHI medium at 37°C. The graphs represent averages from triplicate measurements. The CRISPRi-strains were grown with and without IPTG, as indicated. Note that all CRISPRi strains, in this study, started growing rapidly after approximately 10 hours. This phenomenon is most likely caused by reduced functionality of the CRISPRi system in this experimental setup beyond the 10-hour mark. (**B**) Micrographs of the same strains as in (A) showing phase contrast (PC) and fluorescence microscopy of cells stained with the cell wall label VanFL and the nucleoid label DAPI. CRISPRi strains were grown in medium with IPTG to induce the CRISPRi-system. White arrows point to cells with perturbed septum formation and abnormal nucleoid staining. The scale bars are 2 µm. (**C**) Violin plots of the cell areas (in µm^2^) of JE2 wild-type (1.09 ± 0.22 µm^2^), Δ*cozEa*::Tn (1.01 ± 0.22 µm^2^), Δ*cozEb*::Tn (1.09 ± 0.24 µm^2^), MDB19 (1.25 ± 0.36 µm^2^), and MDB21 (1.31 ± 0.39 µm^2^), determined using MicrobeJ. Significant differences between the strains are indicted with asterisks (* indicates a P-value of < 0.05, ** indicates a P-value of < 0.01, and *** indicates a P-value of < 0.001, derived from a Mann-Whitney test). The number of cells analyzed for each strain is indicated in the figure. (**D**) Schematic outline of the different cell cycle phases used to classify the cells in (E). Cells in phase 1 are non-dividing cells without visible septa, cells in phase 2 are actively dividing cells with incomplete septa, and cells in phase 3 are dividing cells with fully formed septa. (**E**) Frequency of cells in each of the three cell cycle phases for JE2 wild-type, Δ*cozEa*::Tn, Δ*cozEb*::Tn, MDB19, and MDB21. The distributions were obtained by manual counting the different cell cycle phases of 100-150 randomly selected VanFL stained cells from each strain. (**E**) TEM micrographs of uninduced and induced CRISPRi(cozEa+cozEb) (MDB19) cells. White arrows point to cells with aberrant septum formation. The scale bar is 200 nm.

Similar to what was reported previously (26), we were unable to obtain a double Δ*cozEa*Δ*cozEb* mutant in NCTC8325-4 by allelic replacement using the pMAD-vector (29). We therefore used an established two-plasmid CRISPR interference (CRISPRi) system for knockdown of gene expression (26, 30). In this system, dCas9 is expressed from an IPTG-inducible promoter on one plasmid, and the gene-specific sgRNA is constitutively expressed from the other. Indeed, simultaneous knockdown of *cozEa* and *cozEb* in wild-type backgrounds, knockdown of *cozEa* in the Δ*cozEb* backgrounds (Δ*cozEb*, CRISPRi(*cozEa*)) or *vice versa* caused a clear growth reduction in growth in both JE2 and NCTC8325-4 (**Fig. 1A**, **Fig. S1B**, **Fig. S2A**). These cells also exhibited perturbed cell sizes, shapes and septa compared to the wild-type cells and single deletions (**Fig. 1B-C, 1F**, **Fig. S2B-C**). By categorizing the cells into three cell cycle states (**Fig. 1D**), we observed over-representation of phase 1 cells (cells before initiating septum synthesis) and under-representation of phase 2 (cells with incomplete septa) and phase 3 cells (cell with complete septa) (**Fig. 1E**, **Fig. S2D**), showing that coordination of cell division is disturbed in cells lacking both CozE proteins. More specifically, this indicates that initiation of septum synthesis is inhibited in these cells.

### Cells lacking both CozE proteins have mis-localized cell wall synthesis

Localization of peptidoglycan synthesis was further investigated by labelling cells with the fluorescent D-amino acid 7-hydroxycoumarincarbonylamino-D-alanine (HADA) (31). Since HADA is incorporated into newly synthesized peptidoglycan via transpeptidation during the labeling period, it is possible to distinguish sites of active growth. In this study, the cells were incubated with HADA for 2 minutes at 37 °C. Single deletions of *cozEa* or *cozEb* did not have an impact on the localization of nascent peptidoglycan in JE2 nor NCTC8325-4, it was located properly in the septal region (**Fig. 2**). However, when both CozE proteins were absent, the HADA signals were highly heterogeneous with respect to signal intensity, and frequently appeared as clustered aggregates instead of being localized at the septum (**Fig. 2-D**). These observations show that peptidoglycan synthesis in *S. aureus* is disturbed when both CozE proteins are absent.

**Fig. 2.**
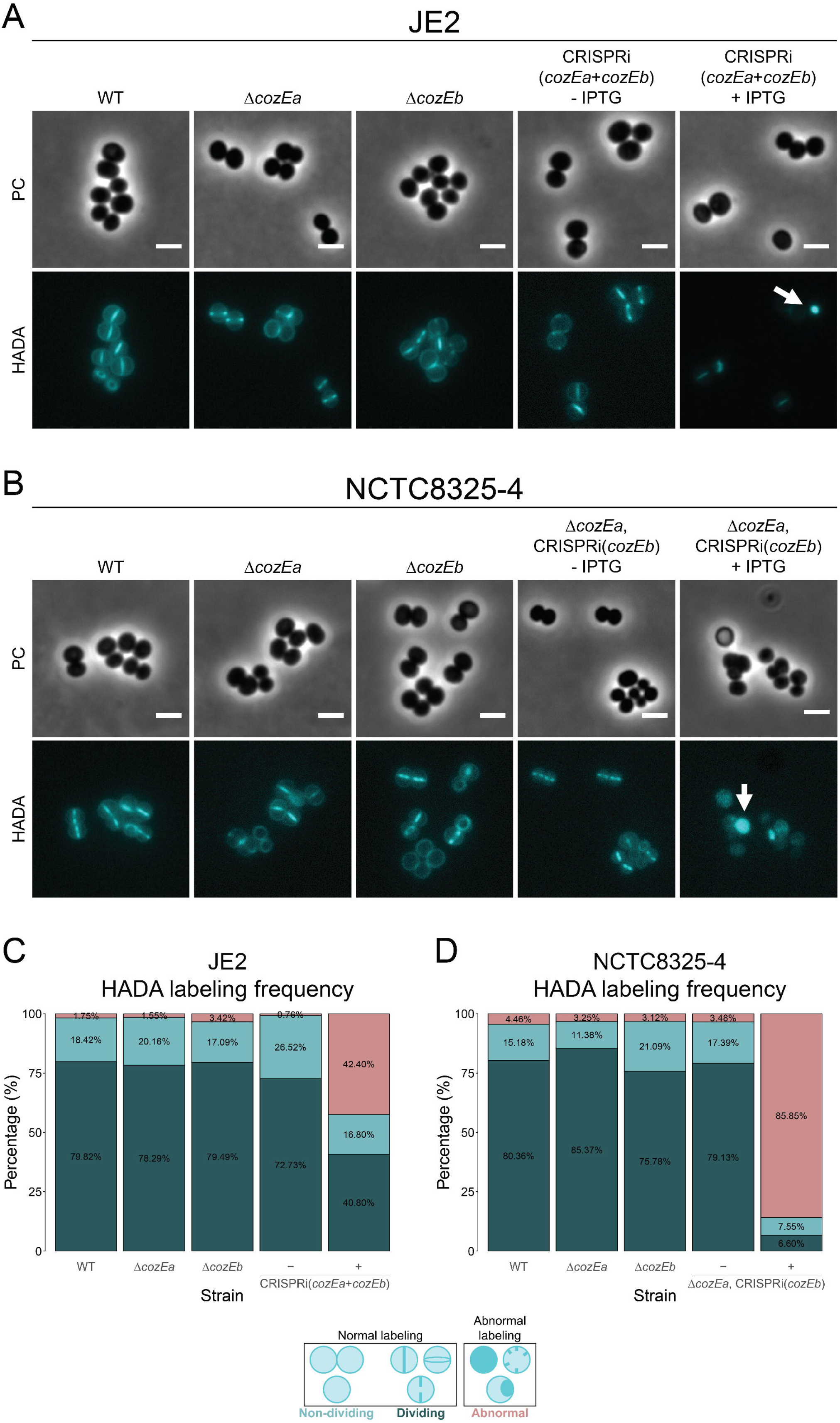
Localization of peptidoglycan synthesis in *S. aureus* JE2 and NCTC8325-4. (**A**) Phase contrast (PC) and fluorescence micrographs of HADA-labeled JE2 wild-type (MDB9), Δ*cozEa* (MDB38), Δ*cozEb* (MDB10), and CRISPRi(*cozEa*+*cozEb*) (MDB19). (**B**) Phase contrast (PC) and fluorescence micrographs of HADA-labeled NCTC8325-4 wild-type (MDB1), Δ*cozEa* (MDB2), Δ*cozEb* (MDB3), and Δ*cozEa*, CRISPRi(*cozEb*) (MDB11). Both JE2 and NCTC8325-4 cells were incubated with HADA for 2 minutes at 37 °C. The double *cozE* mutants (MDB19 and MDB11) were grown with and without IPTG. The induced cells displayed irregular HADA staining, indicated by the white arrows. The scale bars are 2 µm. Frequency of cells with normal (dividing and non-dividing) and abnormal HADA labeling for (**C**) JE2 wild-type (MDB9), Δ*cozEa* (MDB38), Δ*cozEb* (MDB10), and CRISPRi(*cozEa*+*cozEb*) (MDB19), and (**D**) NCTC8325-4 wild-type (MDB1), Δ*cozEa* (MDB2), Δ*cozEb* (MDB3), and Δ*cozEa*, CRISPRi(*cozEb*) (MDB11). The distributions were obtained by manual counting the labeling pattern of 100-150 randomly selected HADA stained cells from each strain.

### The *cozE* genes have a synthetic link to genes involved in LTA synthesis

Mutants lacking both CozE proteins exhibited distinctive phenotypic traits, such as morphological abnormalities and impaired control of septum formation (**Fig. 1**, **Fig. S2**, **Fig. 2**) (25). This resembles the phenotypes of *S. aureus* mutants with defects in LTA biosynthesis (Δ*ltaS*, Δ*ltaA* or Δ*ugtP*) (4, 6, 7, 9, 14) (**Fig. 3A**). In this context it is also interesting to note that in a study by Corrigan et al. (12), re-sequencing of *ltaS* deletion mutants resulted in potential suppressor mutations in *cozEb* (SAOUHSC_01358). This prompted us to investigate a potential link between CozE proteins and LTA synthesis. To screen for potential functional links between *cozE* and LTA synthesis genes, sgRNAs targeting *ltaS* and *ugtP*-*ltaA* were made (**Fig. 3A**, *ugtP* and *ltaA* are in the same operon and are therefore targeted together with CRISPRi). A reduction in growth rate was observed upon depletion of UgtP-LtaA or LtaS in a wild-type background (**Fig. S3A**). LTA synthesis genes were then knocked down in the Δ*cozEa* and Δ*cozEb* genetic backgrounds, to see whether the absence of these genes affected the growth (**Fig. S3A**). No major effects were evident, although the growth reduction observed in cells depleted of LtaS or UgtP-LtaA appeared to be slightly alleviated in both Δ*cozEa* and Δ*cozEb* backgrounds (indicated by the lengths of the red arrows in **Fig. S3A**). Following up on this, the growth and cell size defects observed upon knockdown of LtaS were indeed less in a Δ*cozEb* genetic background, although the observed effect was relatively minor (**Fig. S4**).

**Fig. 3.**
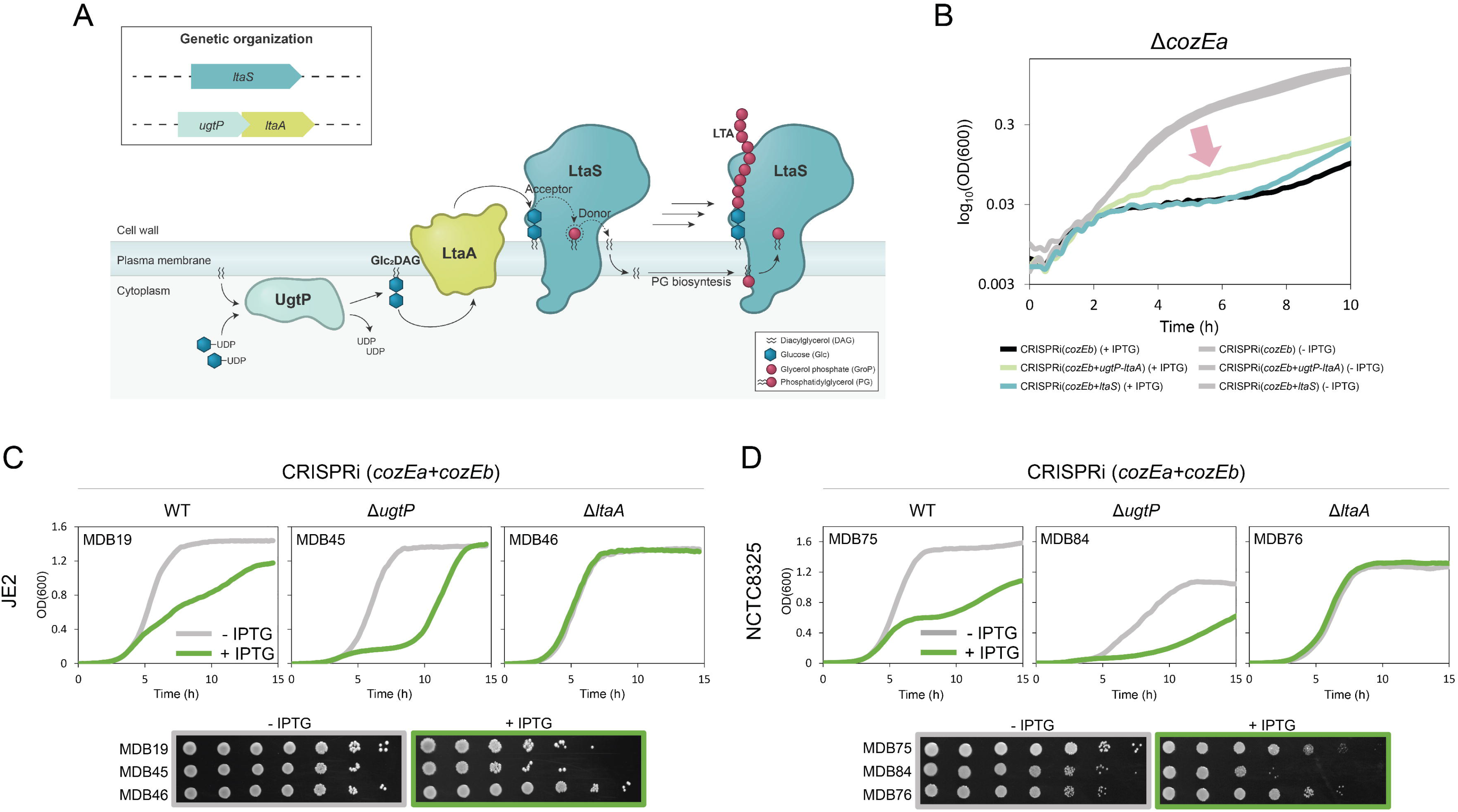
Synthetic genetic relationships between *cozE* genes and genes involved in LTA biosynthesis. (**A**) Schematic overview of the LTA biosynthetic pathway. UgtP (also referred to as YpfP) synthesizes the LTA glycolipid anchor, Glc_2_DAG, from UDP-glucose and diacylglycerol (DAG), which is flipped to the outer membrane leaflet by LtaA. LtaS then synthesizes the LTA polymer by transferring glycerol phosphate units (GroP) derived from phosphatidylglycerol (PG) to the glycolipid anchor. The genetic organization of *ltaS*, *ugtP*, and *ltaA* is indicated in the box. (**B**) The initial 10 hours of growth for MDB11 (Δ*cozEa*, CRISPRi(*cozEb*)), MDB25 (Δ*cozEa*, CRISPRi(*cozEb*+*ugtl*-*ltaA*)), and MDB26 (Δ*cozEa*, CRISPRi(*cozEb*+*ltaS*)) from **Fig. S3** displayed in logarithmic scale to provide a clearer representation of the growth alleviation observed when *ugtP*-*ltaA* was knocked down together with *cozEb* in the Δ*cozEa* background (red arrow). The graphs represent averages from triplicate measurements. The CRISPRi-strains were grown with and without IPTG, as indicated by the colors. **(C-D)** Growth of wild-type, Δ*ugtP*, and Δ*ltaA* cells with double *cozE* knockdown in (**C**) JE2 (MDB19, MDB45, and MDB46) and (**D**) NCTC8325 (MDB75, MDB84, MDB76) in liquid cultures (top panels) and on agar plates (bottom panels). Cells were grown in the presence or absence of IPTG (15 µM for JE2 and 125 µM for NCTC8325) as indicated by the colors. Strain names are indicated in the figure. The graphs represent averages from triplicate measurements.

Subsequently, we proceeded to investigate the growth phenotypes when both CozEa and CozEb were absent simultaneously with the LTA biosynthesis genes (**Fig. 3B**, **Fig. S3B**). Interestingly, when UgtP-LtaA were depleted in a strain lacking both CozE proteins, the growth was improved compared to the control strain only lacking CozE proteins (MDB25 vs MDB11, **Fig. 3B**, **Fig. S3B**), suggesting that the detrimental effect of lacking both CozEa and CozEb is partly alleviated when UgtP and/or LtaA is removed. The same trend, although less clear, was also observed when LtaS was depleted in this background (strain MDB26, **Fig. 3B**, **Fig. S3B**). Together, these observations suggested potential functional links between CozE and glycolipid synthesis and prompted us to investigate the interplay between the *cozE* genes and LTA biosynthesis genes *ugtP* and *ltaA*.

### LtaA and UgtP modulate the essentiality of CozE proteins

To further understand how *ugtP*-*ltaA* knockdown partly alleviated the growth defects of CozEa and CozEb deficient cells, the sgRNAs targeting *cozEa*, *cozEb*, or both *cozEa* and *cozEb* simultaneously, were transformed into *S. aureus* JE2 strains with single deletions of *ugtP* (Δ*ugtP*::Tn) and *ltaA* (Δ*ltaA*::Tn). Both Δ*ugtP* and Δ*ltaA* exhibited growth rates comparable to the wild-type strain, and no alterations in growth were observed upon knockdown of the individual *cozE* genes in the JE2 Δ*ugtP* or Δ*ltaA* cells (**Fig. S5**). Strikingly, however, simultaneous knockdown of *cozEa* and *cozEb* in Δ*ugtP* or Δ*ltaA* background caused dramatic, but opposite, alterations to the growth patterns (partial knockdown in **Fig. 3C** and full knockdown in **Fig. S6A**). In the Δ*ugtP* mutant, depletion of the CozE proteins (MDB45) resulted in a synthetic sick phenotype with reduced growth compared to the cells depleted of only the CozE proteins (MDB19) (**Fig. 3C**, **Fig. S6A**). On the contrary, in the Δ*ltaA* mutant, depletion of CozE proteins (MDB46) had less detrimental effect and the growth patterns were more similar to the wild-type (**Fig. 3C**, **Fig. S6A**), indicating a synthetic viable genetic interaction.

To further confirm the intriguing, opposite growth alterations, we conducted the same analysis in *S. aureus* NCTC8325 cells. For NCTC8325 (**Fig. 3D**, **Fig. S6B**), growth reduction was observed for both Δ*ltaA*, and in particular Δ*ugtP*, compared to the wild-type (**Fig. 3D**). Clearly, however, the same CozE-mediated growth patterns were observed in this strain (**Fig. 3D**). Double depletion of CozEa and CozEb was detrimental for growth in a wild-type background (MDB75), and while the growth was further reduced in Δ*ugtP* (MDB84), this effect was alleviated in Δ*ltaA* (MDB76) (**Fig. 3D, Fig. S6B**), confirming that these genetic links are conserved across strains.

Single-cell analyses of the combined mutants further corroborated the pairwise synthetic genetic interactions between *ugtP*, *ltaA*, and *cozE* (**Fig. 4**, **Fig. S7**). The mis-regulation of septal synthesis and cell size defects previously observed in cells depleted of both CozE proteins, were further elevated in the absence of UgtP, as observed by phase contrast imaging and staining with VanFL (MDB19 and MDB45, **Fig. 4A-B**). Conversely, in the Δ*ltaA* cells, depletion of CozEa and CozEb did not yield the same morphological abnormalities (MDB46, **Fig. 4A**), and the cell size distribution in this mutant more closely resembled that of the control strain (MDB46 and MDB44, **Fig. 4B**). We also stained these cells with DAPI to visualize their nucleoids, as previous observations have demonstrated perturbed DAPI staining in *S. aureus* cells with simultaneous depletion of CozEa and CozEb (**Fig. 1A**, **Fig. S2A**) (26). As expected, the highly irregular and distorted DAPI staining pattern was observed when CozE proteins were depleted in the strain lacking UgtP (MDB19 and MDB45, **Fig. 4A**), but the phenotype was rescued in the Δ*ltaA* background (MDB46 and MDB44, **Fig. 4A**).

**Fig. 4.**
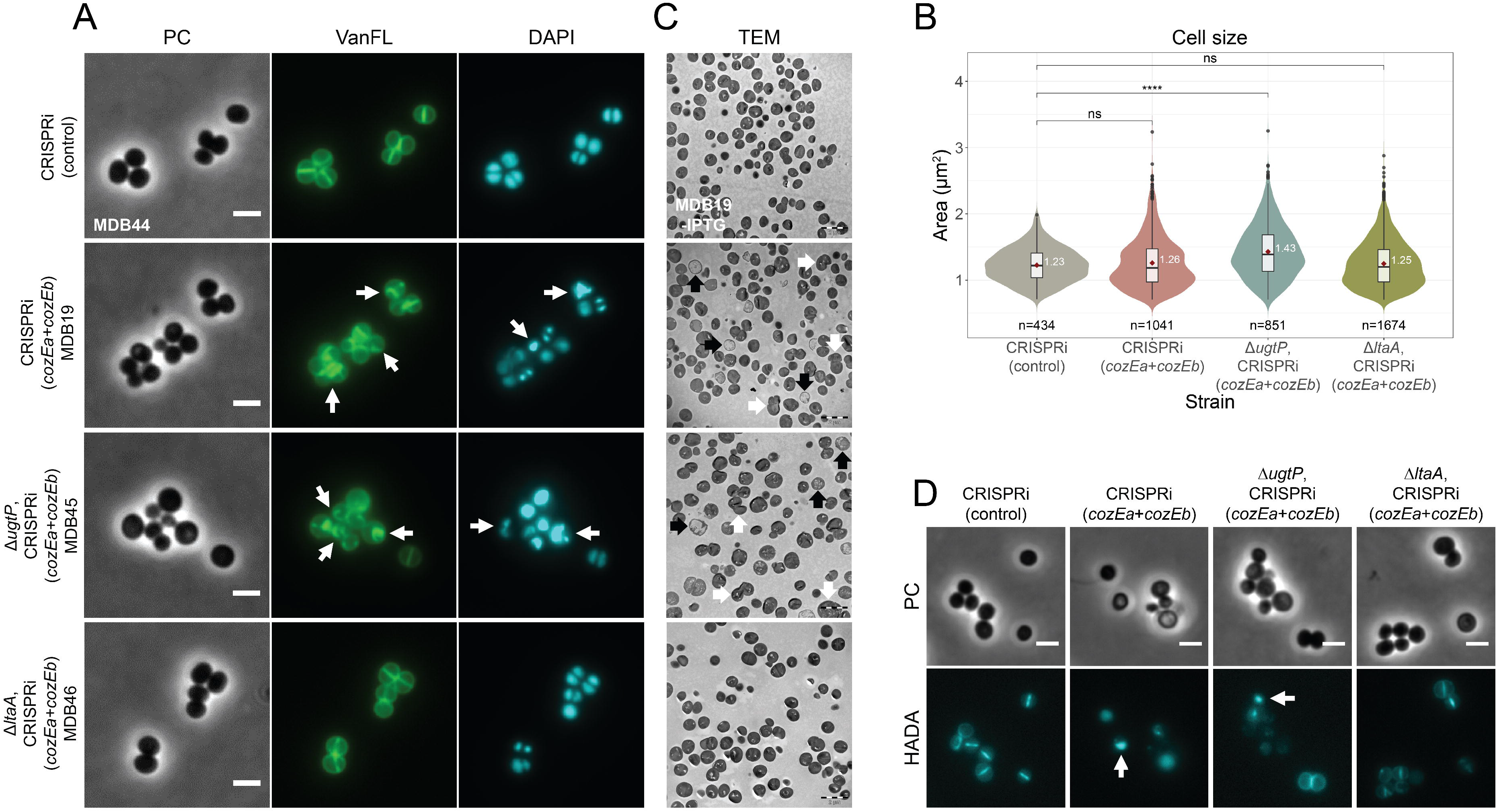
Morphological analysis of CozE depletion in different genetic backgrounds, wild-type, Δ *ugtP*, and Δ *ltaA*, in *S. aureus* JE2. (**A**) Micrographs of JE2 CRISPRi(control) (MDB44), CRISPRi(*cozEa*+*cozEb*) (MDB19), Δ*ugtP*, CRISPRi(*cozEa*+*cozEb*) (MDB45), and Δ*ltaA*, CRISPRi(*cozEa*+*cozEb*) (MDB46) showing phase contrast (PC) and fluorescence microscopy of cells stained with the cell wall label VanFL and the nucleoid label DAPI. The cells were grown in medium with IPTG for induction of the CRISPRi system. White arrows point to cells with perturbed septum formation and abnormal nucleoid staining. The scale bars are 2 µm. (**B**) Violin plots of the cell areas (in µm^2^) of the same cells as in (A), MDB44 (1.23 ± 0.25 µm^2^), MDB19 (1.26 ± 0.38 µm^2^), MDB45 (1.43 ± 0.40 µm^2^), and MDB46 (1.25 ± 0.35 µm^2^), determined using MicrobeJ. Significant differences between the strains are indicted with asterisks (* indicates a P-value of < 0.05, ** indicates a P-value of < 0.01, and *** indicates a P-value of < 0.001, derived from a Mann-Whitney test). The number of cells analyzed for each strain is indicated in the figure. (**C**) TEM micrographs of uninduced and induced CRISPRi(*cozEa*+*cozEb*) (MDB19) cells, induced Δ*ugtP*, CRISPRi(*cozEa*+*cozEb*) (MDB45) cells, and induced Δ*ltaA*, CRISPRi(*cozEa*+*cozEb*) (MDB46) cells. White arrows point to cells with perturbed septum formation, while black arrows point to lysed cells. The scale bars are 2 µm. (**D**) Phase contrast (PC) and HADA staining micrographs of the same strains as in (A) and (B) (MDB44, MDB19, MDB45, and MDB46). The cells were incubated with HADA for 2 minutes at 37 °C. The strains were grown in the presence of IPTG for induction of the CRISPRi system. The induced wildtype and Δ*ugtP* cells displayed irregular HADA staining, indicated by the white arrows. The scale bars are 2 µm.

Finally, we performed transmission electron microscopy (TEM) as well as HADA labelling of the same strains. Consistent with previous findings (26), cells lacking both CozE proteins exhibited a large fraction of lysed cells (black arrows in **Fig. 4C**) in addition to cells with misplaced and abnormal septa (white arrows in **Fig. 4C-D** and **Fig. S7A**), and this phenotype was further exacerbated in the Δ*ugtP* background (**Fig. 4C-D**, **Fig. S7B**). However, in the Δ*ltaA* genetic background, the double *cozE* knockdown had a wild-type like appearance, with few lysed cells and virtually no misplaced septa (**Fig. 4C-D**, **Fig. S7C**). Together, these results show that when UgtP is absent, and Glc_2_DAG are not produced (**Fig. 3A**), CozE proteins become more essential. On the other hand, when LtaA is absent, and thus the flipping of Glc_2_DAG to the outer membrane leaflet is reduced, the CozE proteins seem to be less functionally important.

### CozEb, but not CozEa, affects LTA polymer length

Previous studies have demonstrated that *S. aureus ugtP* and *ltaA* deletion mutants displayed growth defects. This has been attributed to the production of abnormally long LTA polymers formed on an alternative lipid anchor in these mutants, as a result of the loss or reduction in Glc_2_DAG on the extracellular leaflet (14, 16). To determine if CozEa and/or CozEb could influence the LTA polymer in *S. aureus*, the relative lengths of LTA polymers of exponential phase *S. aureus* mutants were analyzed by immunoblotting using an anti-LTA antibody. Notably, the LTA polymers were slightly, but consistently, longer in the Δ*cozEb* mutants compared to the wild-type for both strains JE2 and NCTC8325-4 (**Fig. 5A**, **Fig. S8A**). The LTA size in the Δ*cozEa* mutants, on the other hand, was similar to the wild-type for both NCTC8325-4 and JE2 (**Fig. 5A**, **Fig. S8A**). Indeed, complementation experiments further showed that expression of *cozEb*, but not *cozEa*, could recover the LTA to wild-type lengths in the Δ*cozEb* backgrounds (**Fig. 5B**). We also observed that the LTA size did not increase further in the Δ*cozEb* background when *cozEa* was knocked down (**Fig 5A**). The quantity of LTA polymers produced in the cells was not clearly affected in the *cozE* deletion strains, as indicated by similar band intensities in the immunoblots across more than 10 repeated assays (**Fig. 5**, **Fig. S8**). Together, these results suggest that CozEb has a unique role in modulating the length of LTA polymers in *S. aureus*, although it should be noted that the increase in LTA polymer length in the *cozEb* mutants appear to be less dramatic than in cells lacking UgtP and/or LtaA (**Fig. 5A**). In addition, this shows for the first time, that CozEa and CozEb have distinct functions in *S. aureus*, pointing towards an intricate relationship between the two homologues beyond their redundant, overlapping functions.

**Fig. 5.**
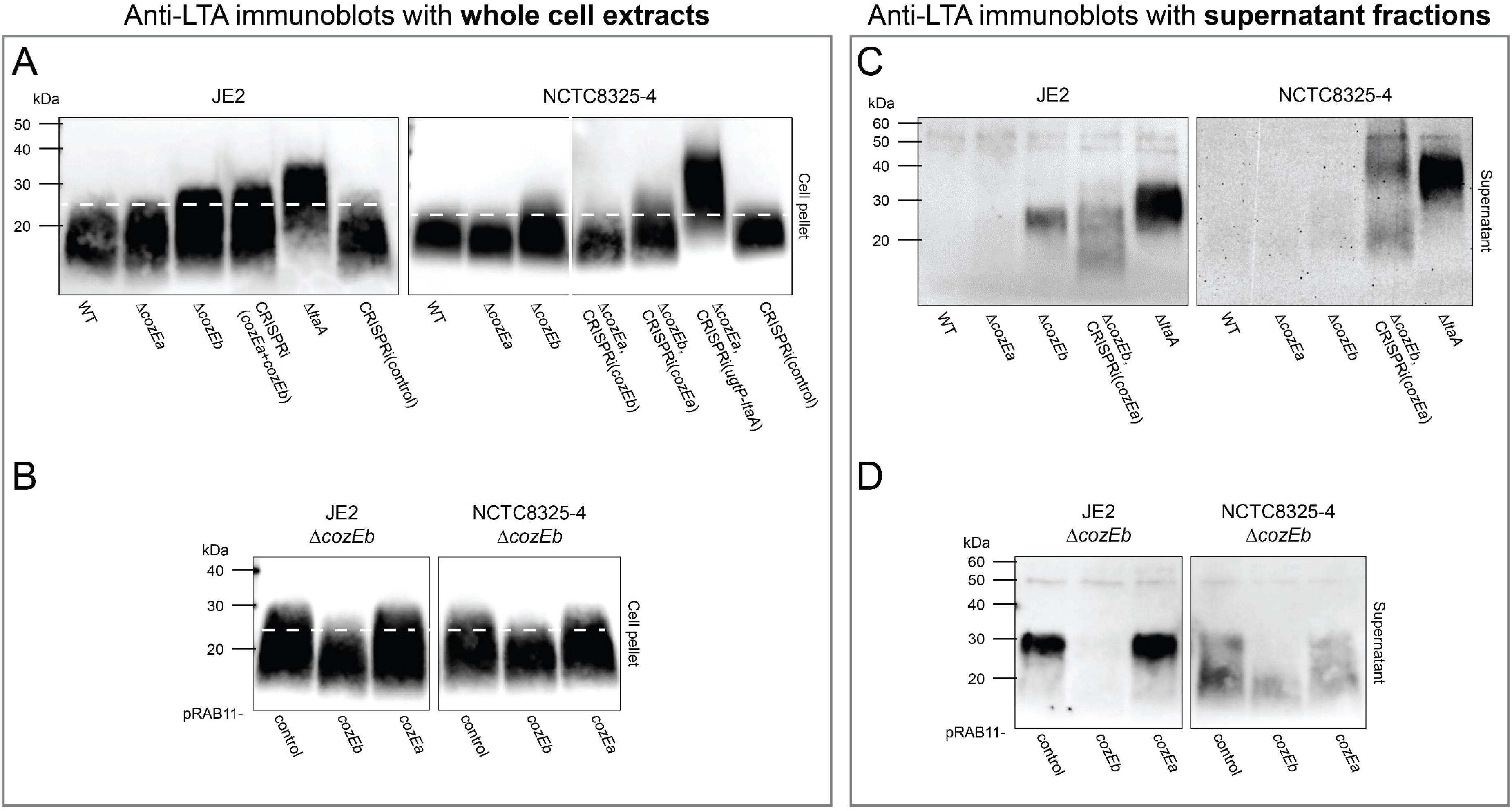
Characterization of LTA polymer length and stability in single and double *cozE* mutants. LTA polymers were detected in whole cell extracts in (A) and (B) and in supernatant fractions in (C) and (D) by immunoblotting with an anti-LTA antibody. (**A**) Immunoblots of wild-type, Δ*cozEa*, Δ*cozEb*, a double *cozE* mutant, a positive control strain, and a CRISPRi-control strain in JE2 (MDB9, MDB38, MDB10, MDB19, MDB40, and MDB44) and NCTC8325-4 (MDB1, MDB2, MDB3, MDB11, MDB12, MDB28, and MM75). (**B**) Complementation experiments in JE2 (MDB61, MDB60, and MDB59) and NCTC8325-4 (MDB63, MDB62, and MDB58). The plasmids pRAB11-*cozEb* and pRAB11-*cozEa*, as well as a pRAB11 control plasmid, were introduced into the Δ*cozEb* mutants. Expression of the plasmid-located genes was induced by addition of 0.004 µg/mL aTc. (A) and (B) illustrate that LTA polymers are consistently longer in the absence of CozEb, and that this phenotype can be complemented with ectopic expression of CozEb but not CozEa. (**C**) Immunoblots of wild-type, Δ*cozEa*, Δ*cozEb*, a double *cozE* mutant, and a positive control strain in JE2 (MDB9, MDB38, MDB10, MDB21, and MDB40) and NCTC8325-4 (MDB1, MDB2, MDB3, MDB12, and MDB69). (**D**) Complementation experiments in JE2 (MDB176, MDB60, and MDB59) and NCTC8325-4 (MDB174, MDB62, and MDB58). The plasmids pRAB11-*cozEb* and pRAB11-*cozEa*, as well as a pRAB11 control plasmid, were introduced into the Δ*cozEb* mutants. Expression of the plasmid-located genes was induced by addition of 0.004 µg/mL aTc. (C) and (D) illustrate that the stability of LTA polymers is compromised in the absence of CozEb, and that it can only be recovered with ectopic expression of CozEb, not CozEa.

### LTA stability is compromised in the absence of CozEb

We also analyzed the presence of LTA in the supernatant fraction of cells lacking CozEa and/or CozEb using the anti-LTA antibody. Detection of LTA in supernatants (which are thus no longer anchored in the cytoplasmic membrane) serves as an indicator of LTA stability. Strikingly, in the JE2 Δ*cozEb* mutant, LTA is clearly being released to the growth medium (**Fig. 5C**, **Fig. S8B**), while no LTA is detected in the supernatant fraction of wild-type or Δ*cozEa*. Although less clear, the same trend is also seen in NCTC8325-4 (**Fig. 5C**, **Fig. S8B**). Furthermore, complementation experiments showed that expression of *cozEb*, but not *cozEa*, could recover the LTA stability in the Δ*cozEb* background (**Fig. 5D**). The size of the released LTA polymers corresponds to the LTA with increased length in the Δ*cozEb* mutant (**Fig. S8C**), suggesting that only the abnormally long LTA polymers are unstable and consequently released into the growth medium. A Δ*ltaA* mutant, which has previously shown to release LTA (32), was included as a control in this experiment, and as expected LTA polymers with increased size were detected in the supernatant of this strain. For the double *cozE* mutants (Δ*cozEb*, CRISPRi(*cozEa*)), LTA with a range of sizes was detected in the supernatant. This observation can probably be attributed to the high degree of lysis observed in these double mutants (**Fig. 4C**), which likely leads to release of LTA to the supernatant. Collectively, these findings suggest that CozEb has a role in modulating the stability, as well as the length, of LTA polymers in *S. aureus*.

### CozE proteins are not critical for the membrane localization of UgtP

Next, we asked how CozE proteins could influence the LTA biosynthetic pathway. First, we studied their subcellular localization in detail using strains with chromosomally integrated *cozEa-gfp* and *cozEb-gfp* fusions in their native loci in NCTC8325-4 (MK1582 and MK1584, respectively, **Fig. S9**). Immunoblotting using an anti-GFP antibody demonstrated that the two fusion proteins (67.03 kDa for CozEa-GFP and 71.93 kDa for CozEb-GFP) have relatively similar expression levels (**Fig. S9A**). The fluorescence microscopy analysis revealed an uneven and spotty localization of both CozE proteins in the membrane, without any septum enrichment, similar to what has been reported for CozEb in *S. pneumoniae* (24) (**Fig. S9B**). Interestingly, time-lapse microscopy revealed that the spotty localization is highly dynamic, indicating that both CozEa-GFP and CozEb-GFP move rapidly around in the cell membrane (see the arrows in **Fig. S9C** and **Movie S1 and S2**). While LtaA has been shown to localize uniformly in the membrane, a somewhat spotty membrane localization was reported for UgtP (19). The mechanism of membrane localization for UgtP, a 391 amino acid-long protein without any predicted transmembrane segments, is not established (19), and we therefore asked whether CozE proteins could be involved in this process. A strain with a chromosomally integrated ectopic copy of *gfp-ugtP* expressed from its native promoter was made, and as expected GFP-UgtP displayed a spotty localization in the membrane (**Fig. S9D**). We subsequently performed single and double knockdown of *cozEa* and *cozEb*, however, membrane-enriched localization of GFP-UgtP was still observed in all of these knockdown strains (**Fig. S9D**). Furthermore, bacterial two-hybrid assays, performed to identify potential protein-protein interactions, did not reveal any direct interaction between UgtP and the CozE proteins (**Fig. S9E**), together indicating that neither CozEa nor CozEb are solely responsible for the cellular localization of UgtP.

### CozE proteins modulate LtaA mediated flipping of Glc_2_-DAG

Our results showed that in the absence of LtaA (implying that the translocation of Glc_2_DAG across the membrane is impaired), the functional relevance of the CozE proteins is reduced. Thus, we asked whether the presence of CozEa and/or CozEb could influence the LtaA-mediated flipping of Glc_2_DAG. To answer this question, we performed *in vitro* flipping assays with LtaA proteoliposomes reconstituted alone (33) or in the presence of CozEa and/or CozEb (**Fig. 6, Fig. S10**). Fluorescently labelled Glc_2_DAG (diglucosyl-diacylglycerol-NBD) was also incorporated in the proteoliposomes as a reporter of flipping activity (**Fig. 6A**). Glc_2_DAG-NBD in the outer leaflet undergo fluorescence quenching upon addition of the membrane-impermeable reducing agent sodium dithionite, and the extent of flipping can therefore be determined by measuring the percentage of fluorescence remaining after sodium dithionite addition (**Fig. 6A**). Our results indicate that CozEb, but not CozEa, slightly decrease LtaA flipping activity *in vitro* (**Fig. 6B**). Interestingly, when CozEa is present, the effect of CozEb is abrogated, thus protecting LtaA from the apparent inhibitory effect of CozEb (**Fig. 6B**).

**Fig. 6.**
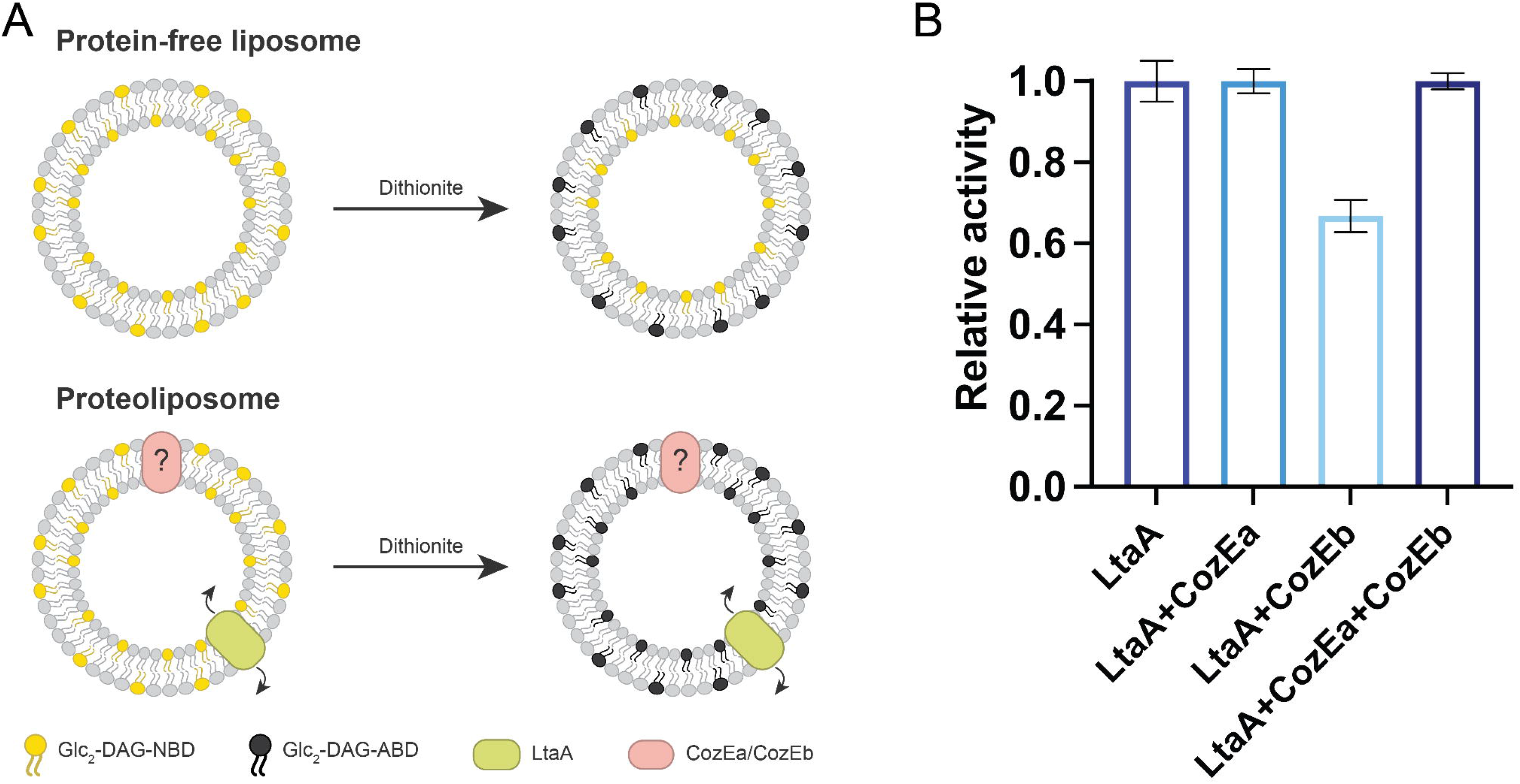
LtaA-catalyzed Glc_2_DAG flipping in the presence of CozE proteins. (**A**) Schematic overview of the *in vitro* flipping assay. Liposomes are reconstituted with NBD-labeled Glc_2_DAG LTA anchors (yellow) that distribute equally in the two membrane leaflets. The fluorescently labeled LTA anchors undergo fluorescence quenching (black) upon addition of sodium dithionite, causing 50% fluorescence loss in protein-free liposomes due to reduction of only the outer-leaflet fluorophores. However, when LtaA (green), known to facilitate the translocation of Glc_2_DAG across the membrane, is present a larger proportion of the fluorophores are reduced due to the flipping activity of LtaA. (**B**) Relative activity of LtaA in the presence of CozE proteins. Relative activity = 100 × (F’ *_i_* − F’ *_liposomes_*)/(F’ *_LtaA_* − F’ *_liposomes_*), where *i* correspond to the fluorescence at the plateau for protein-free liposomes, and F^’^ correspond to the fluorescence at the plateau for proteoliposomes containing (i) LtaA; (ii) LtaA and CozEa (1:1 molar ratio); (iii) LtaA and CozEb (1:1 molar ratio); (iv) LtaA, CozEa, and CozEb (1:1:1 molar ratio). Error bars show + /− s.d. of technical replicates, n = 3.

## Discussion

In line with previous observations in *S. aureus* SH1000 (26), we here show that the two homologous membrane proteins CozEa and CozEb constitute a synthetic lethal gene pair across *S. aureus* strains. Cell cycle analyses of CozE depleted strains show an enrichment of cells without septa (phase 1), and the aberrant localization of cell wall and peptidoglycan synthesis in these cells provides additional validation for their effect on cell division and septum formation in *S. aureus*. The effect of CozE proteins on morphology and growth have consequently directed most studies of these proteins towards their direct interaction with the cell wall and cell division synthesis machineries (21, 24–26, 34). In the present study we have made discoveries that link the function of these proteins to LTA biosynthesis in *S. aureus*. It is indeed well established that mutants of *ugtP*, *ltaA*, and *ltaS* cause cell division defects (4, 8, 14). CozEa/CozEb do not interact with any of the PBPs found in *S. aureus* (26). However, LTA biosynthetic proteins are shown to associate with a number of proteins involved in cell wall synthesis and division, including PBP1, PBP2, PBP3, FtsW, EzrA, DivIB, DivIC, and FtsL (19). Furthermore, it has been demonstrated that LTA affects the activity of cell wall hydrolases required for proper cell splitting (14, 35). It should also be noted that in *B. subtilis*, changes in UgtP activity directly affected the assembly of the division ring, and UgtP has been suggested to function as a metabolic sensor governing cell size in this bacterium (36). In this context, the findings presented in this study suggest that the cell division phenotypes associated with CozE proteins in *S. aureus* may, at least to some extent, be attributed to the effect CozE have on the LTA biosynthetic pathway (26, 37), although other mechanisms, such as their direct interaction with EzrA may also play a role (26).

Our data indeed demonstrate an intricate interplay between CozE proteins and the LTA biosynthetic pathway in *S. aureus*. Firstly, it was noted that the mutants lacking both CozE proteins (**Fig. 1**, **Fig. S2**) were phenotypically similar to staphylococcal cells lacking LTA biosynthetic genes (6, 7, 14). Secondly, our genetic analyses demonstrated that the essentiality of CozE proteins is altered in the absence of LTA biosynthetic genes. Particularly interesting, when the flippase *ltaA* is deleted, resulting in accumulation of the glycolipid anchor Glc_2_DAG on the intracellular membrane leaflet and reduced levels of Glc_2_DAG on the extracellular leaflet, the essentiality of CozE proteins appears to be alleviated (**Fig. 3**, **Fig. 4**). The opposite seems to be true for Δ*ugtP*. In these cells, where the glycolipid anchor is lacking completely, the absence of CozEa and CozEb results in an aggravated growth defect compared to the wild-type background (see further discussion below). In line with these genetic links, suppressor mutations in *cozEb* have previously been reported in a Δ*ltaS* mutant (12). Thirdly, we here observed that deletion of CozEb resulted in slightly longer and more unstable LTA polymers compared to the wild-type (**Fig. 5**). This has previously been associated with cells lacking (Δ*ugtP*) or having reduced levels of (Δ*ltaA*) the LTA anchor Glc_2_DAG on the extracellular leaflet (16). In the absence of Glc_2_DAG, PG can be used as an alternative starter unit for LTA synthesis (6), and LTA polymers formed on PG are longer and more unstable than those formed on Glc_2_DAG (16).

In *S. pneumoniae*, CozE proteins are involved in spatiotemporal localization of peptidoglycan synthesis, probably through their interaction and control of the bifunctional class A PBPs (21, 25). The link between CozE proteins and teichoic acids observed here, may however be relevant also in this species. It has for example been shown that the growth reduction caused by *cozE* knockdown in *S. pneumoniae* is improved in a mutant where the “LTA anchor formation protein B” gene (*lafB*, also named *cpoA*, involved in synthesis of glycolipid Gal-Glc-DAG) is deleted (38). Furthermore, in a *S. pneumoniae tacL* (encoding the lipoteichoic acid ligase required for LTA assembly) deletion mutant, suppressors in *cozE* were found (39).

The effect CozE proteins have on LTA biosynthesis in *S. aureus* may be explained by their effect on membrane homeostasis such as the ability to modulate the flipping activity of LtaA, as demonstrated by *in vitro* assays (**Fig. 6**). The bilayer distribution of Glc_2_DAG in the membrane may be changed in *cozE* mutant cells, thus disturbing the synthesis of LTA polymers, as observed in the Δ*cozEb* background. However, it still remains to be understood whether CozE proteins influence LtaA directly or indirectly and why CozEa and CozEb apparently have opposite functions in this regard. It is also interesting to note that both CozEa and CozEb are highly dynamic membrane proteins, and it could be speculated that their role is important not only for the flipping of lipids, but also for the lateral dynamics of lipids and/or fluidity of the membrane. Also worth noting in this context, is that a Δ*cozEb* mutant in *S. pneumoniae* has been shown to display increased susceptibility to the membrane targeting antibiotic daptomycin (34, 40, 41). Furthermore, our results showing that the essentiality of CozE proteins are oppositely affected in a Δ*ltaA* and Δ*ugtP* backgrounds (**Fig. 3**), suggest that CozE proteins may be involved in the bilayer distribution of lipids. I.e., accumulation of Glc_2_DAG (in Δ*ltaA*) somehow alleviates the need for CozE proteins observed in the wild-type (which probably has a more even distribution of Glc_2_DAG) and particularly in Δ*ugtP* (which do not contain Glc_2_DAG). Lipid scramblases are known to play important functions in regulating lipid distributions in eukaryotic cells (42). It is tempting to speculate that CozE proteins function as bacterial membrane scramblases, and in that way affect many membrane-associated processes including peptidoglycan and teichoic acid biosynthesis. However, this hypothesis needs further investigation, for example by structural analysis of potential interactions between the highly dynamic CozE proteins and different lipids and/or by analyzing the distribution of lipids between the bilayers in different genetic backgrounds.

Our results also demonstrate, for the first time, that CozEa and CozEb are not fully redundant in *S. aureus*, as deletion of *cozEb*, but not *cozEa*, resulted in longer and more unstable LTA polymers (**Fig. 5**). This is not entirely surprising given the different roles of the two CozE paralogs in *S. pneumoniae* (25). The notion of the unique functionality of these paralogs is indeed supported by a phylogenetic analysis of CozE proteins from *Streptococcaceae* (25) and *Staphylococcaceae* (**Fig. S11**). The phylogenetic analysis of CozE from 28 different species within the *Staphylococcaceae* family demonstrated that each species encodes two CozE proteins, and that the two paralogs cluster into two separate subgroups, corresponding to CozEa and CozEb (blue and red, respectively in **Fig. S11**) for the genera *Staphylococcus* and *Macrococcus*. It should be noted that CozE proteins from more distantly related genera (*Jeotgalicoccus*, *Salinicoccus*, and *Nosocomiicoccus*) do not display this subclassification, but instead cluster into a separate group that is phylogenetically closer to CozEa (green in **Fig. S11**), indicating that the function of CozEb may be unique for *Staphylococcus* and *Macrococcus*.

It has already been shown that CozEb can act as target for antibody-based infection treatment in *S. pneumoniae* (34). Functional insights into the role of CozE proteins in different bacteria are needed to further explore their potential in anti-microbial or anti-infection treatment. The novel functions of CozE proteins demonstrated here, reveal that these proteins are new players in the control of LTA biosynthesis and membrane homeostasis in *S. aureus*. Future work should aim at further deciphering the interplay between the CozE proteins and between CozE proteins and LTA synthesis, membrane homeostasis, and cell division in *S. aureus* and other bacteria.

## Methods

### Bacterial strains and growth conditions

Strains used in this work are listed in **Table S1**. *S. aureus* strains NCTC8325-4, NCTC8345, and JE2-USA300 (called JE2 here) were grown in BHI medium with shaking or on BHI agar plates at 37°C, if not stated otherwise. *E. coli* strains IM08B, XL1-Blue, and BTH101 were grown in LB medium with shaking or on LA plates at 37°C, if not stated otherwise. When appropriate, antibiotics were added for selection: 100 µg/mL ampicillin and/or 50 µg/mL kanamycin for *E. coli*, 100 or 1000 µg/mL spectinomycin (for NCTC8325-4 and JE2, respectively), 5 µg/mL erythromycin and/or 10 µg/mL chloramphenicol for *S. aureus*. Isopropyl β-D-1-thiogalactopyranoside (IPTG) and anhydrotetracycline (aTc) were added for induction of transcription when needed.

For transformation of *E. coli*, chemically competent IM08B cells were prepared using calcium-chloride treatment followed by transformation with heat shock according to standard protocols. *S. aureus* strains were transformed by electroporation with plasmids isolated from *E. coli* IM08B, as described previously (43).

### Strain construction

All strains used in this work are listed in **Table S1**, plasmids are listed in **Table S2**, while primers used for the cloning are listed in **Table S3**. Every construct was verified by PCR and sequencing.

#### Deletion of cozEa and cozEb (ΔcozEa::spc and ΔcozEb::spc)

Deletion of *cozEa* or *cozEb* in *S. aureus* NCTC8325-4 was achieved using the temperature-sensitive pMAD system, following the same approach as described before (26).

#### Deletion of *ugtP* (Δ*ugtP::spc*)

Deletion of *ugtP* in *S. aureus* NCTC8325 was achieved using the temperature-sensitive pMAD system. To construct pMAD-Δ*ugtP*::*spc*, three DNA fragments were initially amplified: (1) the *ugtP* upstream sequence (“ugtP_up”), (2) a spectinomycin resistance cassette (“spc”), and (3) the *ugtP* downstream sequence (“ugtP_down”), using primers listed in **Table S3**. gDNA from *S. aureus* NCTC8325-4 served as template DNA for amplification of both ugtP_up and ugtP_down, while the pCN55 plasmid (44) was used as the template for amplification of *spc*. The primers were designed with overlapping sequences, enabling fusion of the three fragments by overlap extension PCR. The resulting fragment was digested with BamHI (introduced with the outer primer mk506) and NcoI (naturally occurring near the 5’ end of the fragment) and subsequently ligated into the corresponding sites of pMAD. The generated plasmid was verified by PCR and sequencing, and the standard pMAD protocol (29) was used to replace *ugtP* with the *spc*-marker. Note that the *ugtP*-fragment was amplified without any terminator sequence to avoid any downstream effects on the transcription of *ltaA*.

#### Construction of CRISPR interference strains

For knockdown of genes, the two-plasmid CRISPR interference system described previously (26, 30) was used. In this system, the appropriate strains are transformed with a plasmid carrying an IPTG-inducible *dcas9* (pLOW-*dcas9*) and another plasmid carrying a gene-specific sgRNA with constitutive expression (pCG248-sgRNA(*xxx*) or pVL2336-sgRNA(*xxx*), where *xxx* denotes the target gene). The sgRNA plasmids were constructed using inverse PCR in pCG248 (26) or Golden Gate cloning in pVL2336 (30), using oligos listed in **Table S3**, and verified by PCR and sequencing.

#### Construction of pRAB11 plasmids used for complementation

The genes of *cozEa* and *cozEb* were initially amplified from *S. aureus* SH1000 gDNA with primers containing KpnI and EcoRI restriction sites as overhangs (see **Table S3**). Purified PCR products and the plasmid pRAB11 (45) were digested with KpnI and EcoRI, and subsequently ligated using T4 DNA Ligase. Ligation mixtures were transformed into *E. coli* IM08B, and the plasmids were verified by PCR and sequencing before being electroporated into *S. aureus*.

#### Construction of chromosomally integrated *cozEa-gfp* and *cozEb-gfp* fusions

The temperature-sensitive pMAD system was used to GFP-tag *cozEa* and *cozEb* in their native loci in *S. aureus* NCTC8325-4. For construction of pMAD-*cozEa-m(sf)gfp_spc*, the *cozEa-gfp* fusion was amplified from plasmid pLOW-*cozEa*-*m(sf)gfp* using primers mk432 and mk433, while the spectinomycin resistance cassette spliced with the *cozEa* downstream region was amplified from plasmid pMAD-*cozEa*::*spc* using primers mk188 and mk434. The two fragments were fused by overlap extension PCR and ligated into pMAD using the NcoI and SalI restriction sites introduced with the primers. Similarly, for pMAD-*cozEb-m(sf)gfp_spc*, the *cozEb-gfp* fusion was amplified from plasmid pLOW-*cozEb-m(sf)gfp* using primers mk435 and mk433, while the *spc* cassette spliced with the *cozEb* downstream region was amplified from plasmid pMAD-*cozEb::spc* using primers mk188 and mk436. These two fragments were also fused by overlap extension PCR and ligated into pMAD using the NcoI and SalI restriction sites introduced with the primers. Finally, a standard pMAD protocol (29) was used for chromosomal integration of the generated fusions.

#### Construction of a chromosomally integrated *gfp-ugtP* fusion

A *gfp-ugtP* fusion gene, driven by the *ugtP*-promoter, was integrated into a neutral locus (between genes SAOUHSC_03046 and SAOUHSC_03047) on the *S. aureus* NCTC8325-4 chromosome using the temperature-sensitive pMAD system. To construct the plasmid pMAD-P_ugtP_*-m(sf)gfp-ugtP*_*spc*, *gfp* was first fused to the 5’ end of *ugtP* by restriction cloning. *ugtP* was amplified using gDNA from *S. aureus* NCTC8325-4 as template and ligated into the NcoI and BamHI restriction sites of pLOW-*m(sf)gfp*-SA1477 to produce plasmid pLOW-*m(sf)gfp-ugtP*. The four fragments constituting the insert of the pMAD-P_ugtP_*-m(sf)gfp*-*ugtP*_*spc* plasmids were then amplified: (1) the upstream integration region (“ori_up”), (2) the *ugtP*-promoter (“P_ugtP_”), (3) the *gfp*-*ugtP* fusion gene (“gfp-ugtP”), and (4) a spectinomycin resistance cassette spliced with the DNA sequence of the downstream integration region (“*spc*+ori_down”). Both ori_up and P_ugtP_ were amplified using gDNA from *S. aureus* NCTC8325-4 as template, while purified pLOW-*m(sf)gfp*-*ugtP* and pMAD-ori-*parS* were used as template DNA for amplification of gfp-ugtP and spc+ori_down, respectively. All primers used for the forementioned amplifications are listed in **Table S3**. The four fragments were subsequently spliced by overlap extension PCR and ligated into pMAD, using the EcoRI and SalI restriction sites introduced with the outer primers. Finally, a standard pMAD protocol (29) was used to integrate the generated fusion into the chromosome of *S. aureus* NCTC8325-4.

#### Construction of pET19b-cozEa and pET19b-cozEb

*cozEa* and *cozEb* were amplified with the primer pairs mk508/mk509 and mk510/mk512, respectively, producing fragments with flanking SpeI and BamHI restriction sites introduced in the primers. The vector LtaA-pET19b and the fragments, were digested with SpeI and BamHI, and the digested fragments were ligated into the vector and transformed into *E. coli*. The constructs were verified by PCR and sequencing.

### Growth assays in liquid media

To measure growth in liquid medium, the bacterial strains to be monitored were initially grown overnight in BHI medium with the respective antibiotics. They were then diluted 1:1000 in fresh BHI medium supplemented with the respective antibiotics and inducers, when appropriate. The bacterial dilutions were applied to a 96-well microtiter plate and incubated in a plate reader at 37°C for 18-20 hours. OD_600_ measurements were taken every 10 minutes, with a brief shaking of the plate for 2-5 seconds before each measurement. All growth curves in this work are the mean value of three replicate measurements.

### Spotting assays

To assess growth on solid medium, cells grown overnight in BHI medium were serially 10-fold diluted in fresh BHI medium with antibiotics and IPTG for induction, when appropriate. Each overnight culture and its serial dilutions were spotted onto the appropriate BHI agar plates with a volume of 2 µL. The plates were incubated aerobically at 37°C for 17 to 20 h. Images of the plates were captured using a Gel Doc™ XR+ Imager (Bio-Rad Laboratories).

### Epifluorescence- and phase contrast microscopy

For microscopy analyses, strains were firstly grown overnight in BHI medium with the respective antibiotics. The overnight cultures were diluted 1:1000 in fresh BHI medium containing the proper antibiotics and inducers, when appropriate, and incubated until their OD_600_ reached approximately 0.4. In some cases, the cells were stained with fluorescent vancomycin (VanFL, in which a BODIPY fluorophore is linked to a vancomycin molecule (Invitrogen)) and/or DAPI (Invitrogen), at final concentrations of 0.8 µg/mL and 7.5 µg/mL, respectively. In other cases, the cells were stained with a fluorescent D-alanine analogue, HADA (31), at a final concentration of 250 µM. The cultures containing HADA were incubated at 37°C for 2 minutes, before being immediately put on ice to stop bacterial growth. The cells were lastly washed with PBS buffer to remove excess of unbound dye. Bacterial cells were immobilized on agarose pads (1.2 %) before imaging on a Zeiss Axio Observer microscope with ZEN Blue software. The bacteria were visualized with a 100× phase contrast objective, and images were captured using an ORCA-Flash4.0 V3 Digital CMOS camera (Hamamatsu Photonics K.K.). Time-lapse (TL) images of cells expressing GFP-fusions were acquired every third second for 27 seconds using the forementioned equipment.

The distribution of cell sizes among different *S. aureus* strains was determined using MicrobeJ (46). Particles of the strain to be analyzed were detected using a stack of phase contrast images of the given strain in MicrobeJ, every image were subsequently corrected manually by discarding and/or adding cells that were incorrectly detected. In addition to analyzing cell sizes, the cell cycle phases of the bacteria were also analyzed by manual counting the different cell phases (phase 1, 2, or 3) of 100-150 randomly selected VanFL stained cells from each strain. Cell wall synthesis was also analyzed by manual counting the HADA labeling patterns (non-dividing, dividing or abnormal) of 100 of 100-150 randomly selected cells from each strain.

### Transmission electron microscopy

The bacterial strains to be visualized by TEM were first grown overnight in BHI medium with the respective antibiotics and then diluted 1:1000 in fresh BHI with antibiotics and IPTG added when necessary. The diluted bacterial cultures were incubated at 37°C until they reached an OD_600_ of 0.3. Each of the bacterial cultures (10 ml) were carefully mixed with 10 mL fixation solution, containing 2 % (v/v) paraformaldehyde, 0.1 M cacodylate (CaCo) buffer and 1.25 % (v/v) glutaraldehyde solution (grade I). The fixation mixtures were incubated at room temperature for 1 hour, followed by incubation at 4°C overnight. The next day, the cells were centrifugated at 5000 × *g* at 4°C for 5 min, and subsequently washed three times with PBS, pH 7.4, and three times with a 0.1 M CaCo buffer. The cells were then post-fixed for 1 hour with 1 % OsO_4_ in 0.1 M CaCo. The CaCo-washing steps were repeated prior to dehydration, which involved 10 min incubation steps at increasing concentrations of ethanol (70 %, 90 %, 96 %, and 100 %). The samples were next infiltrated with LR White resin by multiple incubation steps with an increasing concentration of the embedding media (mixed with EtOH). First, overnight with a 1:3 ratio of LR White to EtOH, second; approximately 4 hours with a 1:1 ratio, third; 4 hours with a 3:1 ratio, and finally overnight with 100% LR White. The samples were then embedded in 100 % LR White overnight at 60°C by polymerizing the embedding media into a hard block. All sample blocks were sectioned, 60 nm thin, and stained with uranyl acetate and potassium permanganate. A FEI Morgagni™ 268 Transmission electron microscope were used to analyze the samples. Images of the bacteria were captured using a Veleta CCD camera (Olympus Corporation) with an exposure time of ∼1000 ms.

### Immunoblot analysis of lipoteichoic acid in whole cell extracts and supernatants

Overnight cultures were diluted 1:1000 in TSB medium with antibiotics and IPTG for induction, when appropriate, and incubated at 37°C until they reached an OD_600_ between 0.6 and 0.8. The cultures were normalized to an OD_600_ of 0.6 and then harvested by centrifugation at 5400 × *g* for 3 min at 4°C. Detection of LTA was done essentially as described before (14, 32). The supernatants from each strain were transferred to clean tubes, separating them from the pellets, for individual analysis. For pellet fraction analysis, the pellets were resuspended in 50 µL lysis buffer, containing 50 mM Tris-HCl pH 7.4, 150 mM sodium chloride, and 200 µg/mL lysostaphin, before incubation at 37°C for 10 min. The suspensions were then added 50 µL 4× SDS loading buffer and boiled at 95°C for 30 min. The cell lysates were subsequently centrifuged at 16 000 × *g* for 10 min to pellet cellular debris. The supernatants (60 µL) were transferred to clean tubes containing 60 µL dH_2_O. The diluted suspensions were lastly treated with 0.5 µL proteinase K (20 mg/ml) for 2 hours at 50°C. For supernatant fraction analysis, the supernatants were centrifuged at 16 000 × *g* for 10 min, 75 µL of each supernatant were mixed with 25 µL 4× SDS loading buffer and boiled at 95°C for 30 min. They were subsequently centrifuged at 16 000 × *g* for 10 min, and the supernatants (60 µL) were lastly transferred to clean tubes.

The pellet and supernatant samples were separated with SDS-PAGE using a 4-20% Mini-PROTEAN TGX acrylamide gel (Bio-Rad). Next, blotting onto a PVDF membrane was performed using a Trans-Blot Turbo System (Bio-Rad). Afterwards, the membrane was blocked in a PBST solution containing 5 % skimmed milk powder (w/v) for 1 hour at room temperature. After washing with PBST, the membrane was then incubated for 1 hour with an anti-LTA primary antibody (Hycult) (diluted 1:4000 in PBST). Next, the membrane was washed 3 times with PBST to remove unbound antibodies, and then incubated for another hour with an anti-Mouse IgG HRP-conjugate secondary antibody (Promega) (diluted 1:10 000 in PBST). After incubation, unbound antibodies were once again removed by washing the membrane 3 times with PBST. Finally, the membrane was developed using the SuperSignal™ West Pico PLUS Chemiluminescent Substrate kit (Thermo Fisher Scientific), and blot images were captured with an Azure Imager c400 (Azure Biosystems).

### Western blot analysis of the relative expression of GFP-tagged CozEa and CozEb

Overnight cultures of MK1582 (with a *cozEa*-*gfp* fusion) and MK1584 (with a *cozEb*-*gfp* fusion) were diluted 1:100 in TSB medium with 100 µg/mL spectinomycin and incubated at 37°C until they reached an OD_600_ of approximately 0.4. The cultures were normalized to an OD_600_ of 0.4, and subsequently harvested by centrifugation at 4000 × *g* for 1 min at 4°C. The pellets were resuspended in 500 µL TSB buffer before being lysed mechanically using the Fast Prep method with ≤106 µm glass beads at 6 m/s. Insoluble material was removed by centrifugation at 20,000 × g for 2 min. Next, the supernatants were mixed with equal volume 2x SDS loading buffer and boiled at 95°C for 5 min.

The samples were separated with SDS-PAGE using a polyacrylamide gel, which consisted of a 12% separation gel with a 4% stacking gel layered on top. The subsequent blotting and GFP detection steps were carried out as described in “Immunoblot analysis of lipoteichoic acid in whole cell extracts and supernatants”, with the only difference being the selection of antibodies. For detection of GFP-tagged CozE, an anti-GFP primary antibody (Invitrogen) (diluted 1:4000 in PBST) and an anti-Rabbit IgG HRP-conjugated secondary antibody (Promega) (diluted 1:5000 in PBST) were used.

### Bacterial two-hybrid assays

The construction of plasmids and the procedure for bacterial two-hybrid assays were performed as previously described (26). Primers used for plasmid construction are listed in **Table S3**. Briefly, the assay utilizes adenylate cyclase from *Bordetella pertussis* to detect possible protein-protein interactions by making gene fusions of selected genes to the T18 or T25 domains of adenylate cyclase (47) in the plasmid vectors pKT25, pKNT25, pUT18, or pUT18C (Euromedex). The ligated plasmids were transformed into *E. coli* XL1-Blue cells.

Next, isolated plasmids containing fusion genes of opposite domain (one with T18 and another with T25) were co-transformed into *E. coli* BTH101 cells, with kanamycin and ampicillin as selection markers. Five colonies were picked, grown to visible growth in LB medium, and spotted on LA plates with the selection markers, in addition to 40 µg/mL X-gal and 0.5 mM IPTG. After incubation at 30°C protected from the light for 20-48 hours, plates were inspected, positive interactions are indicated by blue colonies, while white colonies indicate the opposite. The presented BATCH assays are representative of 5 independent replicates.

### Phylogenetic analysis

CozE homologues were identified with NCBI BLASTp, using the CozEa and CozEb protein sequence of *S. aureus* NCTC8325 as the queries against species within the *Staphylococcaceae* family. 56 CozE homologues belonging to the *Staphylococcaceae*, including CozEa and CozEb found in *S. aureus*, were selected, and subsequently aligned using Clustal Omega (48). Using IQ-TREE (49), the sequence alignment was then used to construct a maximum likelihood phylogenetic tree. Finally, the phylogenetic tree was visualized and annotated with the Interactive Tree Of Life (iTOL) online tool (50).

### Expression and purification of LtaA, CozEa, and CozEb

LtaA and CozE proteins were expressed and purified using the same protocol previously used for purification of LtaA (33). Briefly, proteins carrying an N-terminal histidine tag were overexpressed in *E. coli* BL21-Gold (DE3) (Stratagene) cells. Cells were grown at 37 °C in Terrific Broth medium supplemented with 1% glucose (wt/vol) and induced with 0.2 mM IPTG. Cells were disrupted and membranes were collected by ultracentrifugation. Membranes were solubilized in 50 mM Tris-HCl, pH 8.0; 200 mM NaCl; 20 mM Imidazole; 15% glycerol (vol/vol); 5 mM β-mercaptoethanol; 1% lauryl maltose neopentyl glycol (wt/vol) (LMNG, Anatrace); 1% N-dodecyl-β-d-maltopyranoside (wt/vol) (DDM, Anatrace) for 2 h at 4 °C. After centrifugation, the proteins were purified by affinity chromatography with Ni-NTA superflow affinity column (Qiagen) as previously described (33).

### Formation of proteoliposomes and *in vitro* flipping assay

LtaA and CozE proteins were reconstituted in unilamellar liposomes as described before (33). Briefly, proteoliposomes were prepared by extrusion through polycarbonate filters (400-nm pore size) from a 3:1 (w/w) mixture of *E. coli* polar lipids and L-α-phosphatidylcholine (Avanti polar lipids) resuspended in 20 mM Tris-HCl pH 8.0; 150 mM NaCl, and 2mM β-mercaptoethanol. After removal of detergent with BioBeads (BioRad), proteoliposomes were centrifugated, washed, and resuspended to a final concentration of 20 mg/ml lipids and flash-frozen in liquid nitrogen and stored at −80 °C until further use. Before performing flipping assays, proteoliposomes were thawed, their resuspension buffer was exchanged to 20 mM MES pH 6.5; 150mM NaCl, and the product of the Glc_2_-DAG-NBD synthesis was incorporated by performing freeze/thaw cycles. Proteoliposomes and protein-free liposomes were diluted to a concentration of 2 mg/ml lipids followed by extrusion through polycarbonate filters (400-nm pore size). Proteoliposomes were immediately used for flipping assays (33). Flipping of Glc_2_-DAG-NBD was assessed by determining the percentage of NBD-fluorescence that is quenched after the addition of a 5 mM sodium dithionite (Sigma) after 200 seconds of starting fluorescence recording. Before finishing data recording, 0.5% Triton X100 was added to permeabilize the liposomes, making all Glc_2_-DAG-NBD molecules accessible to dithionite reduction. The fluorescence after Triton X100 addition was used for baseline calculations. Fluorescence was recorded at 20 °C using a Jasco Fluorimeter.

The excitation and emission wavelengths were 470 and 535 nm, respectively. For analysis, the fluorescence intensity was normalized to F/Fmax. Relative flipping activities were calculated as follows: relative activity = 100 × ((F/F_max_)_i_ − (F/F_max_)_liposomes_)/((F/F_max_)_wt_ − (F/F_max_)_liposomes_), where *i* corresponds to each respective treatment/mutants, *liposomes* corresponds to liposomes without protein, *wt* corresponds to wild-type LtaA, and F/Fmax values correspond to the normalized fluorescence values at the plateau after addition of sodium dithionite. Curves were plotted using GraphPad Prism 8. Time courses of the dithionite-induced fluorescence decay in liposomes were repeated at least three times for each individual experiment.

## Acknowledgments

We would like to thank Marita Torrissen Mårli for help with strain construction, and the NMBU Imaging Center for help with transmission electron microscopy. This work was supported by a grant from the Research Council of Norway (296906, 250976) to MK, from the Agence Nationale de la Recherche (ANR-19-CE15-0011-01) to CG, and from the Swiss National Science Foundation (310030_207974 and PP00P3_198903) to CP.

