## Supplementary figures and images for "The function of CozE proteins is linked to lipoteichoic acid biosynthesis in *Staphylococcus aureus*"

### Movie S1

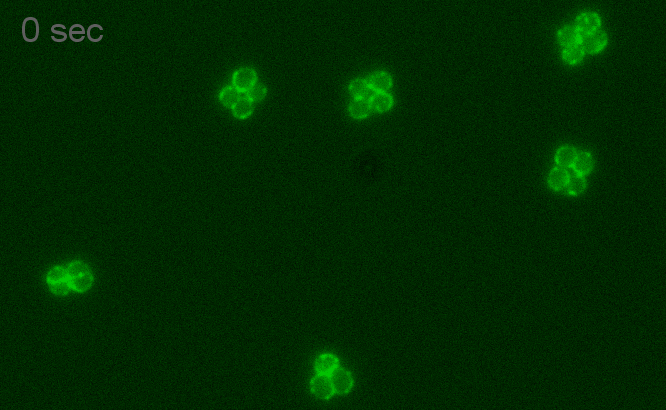

### Movie S2

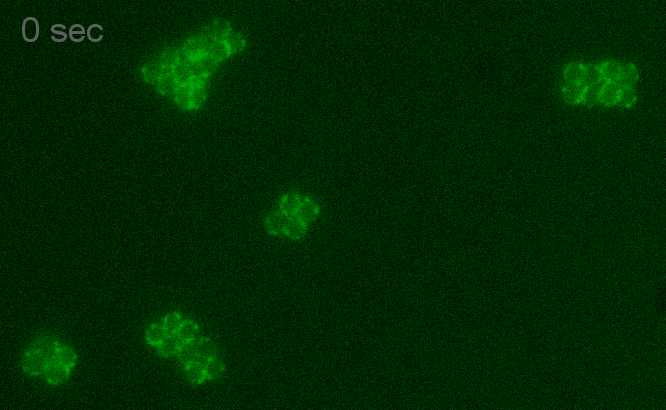
