## Supporting information for "The function of CozE proteins is linked to lipoteichoic acid biosynthesis in *Staphylococcus aureus*"

### Supplementary figures

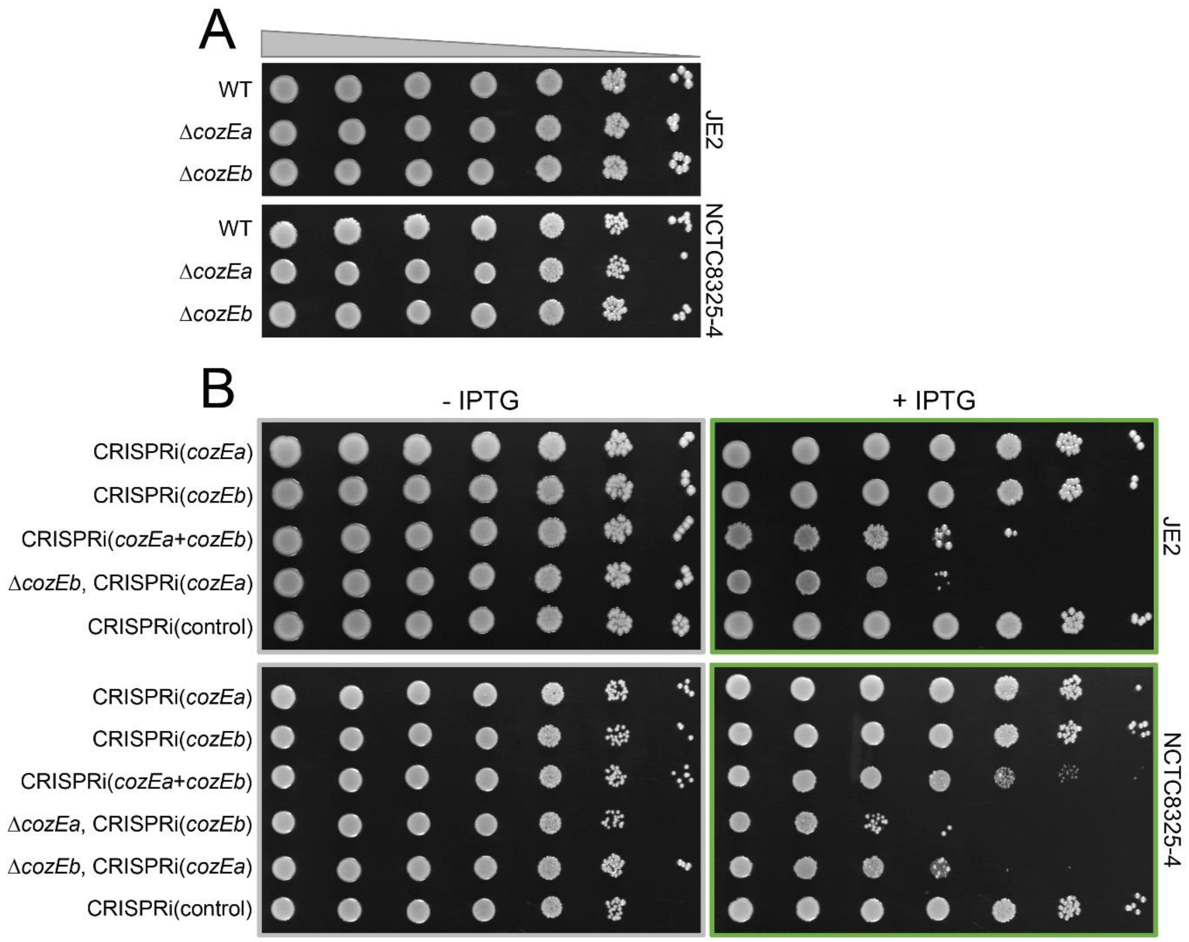


**Fig. S1. Growth of single and double *cozE* mutants in *S. aureus* JE2 and NCTC8325-4.**
(**A**) Growth on solid medium of wild-type, Δ*cozEa*, and Δ*cozEb* in *S. aureus* JE2 (MDB37, MDB38, and MDB10) and NCTC8325-4 (MDB1, MDB2, and MDB3). 10-fold dilution series, made from overnight cultures, were spotted onto agar plates. (**B**) Growth on solid medium of single and double *cozE* knockdown strains in *S. aureus* JE2 (MDB17, MDB18, MDB19, MDB21, and MDB44) and NCTC8325-4 (MDB14, MDB15, MDB13, MDB11, MDB12, and MM75). 10-fold dilution series, made from noninduced overnight cultures, were spotted onto agar plates with and without IPTG, as indicated. Strains carrying a non-targeting sgRNA were used as controls.

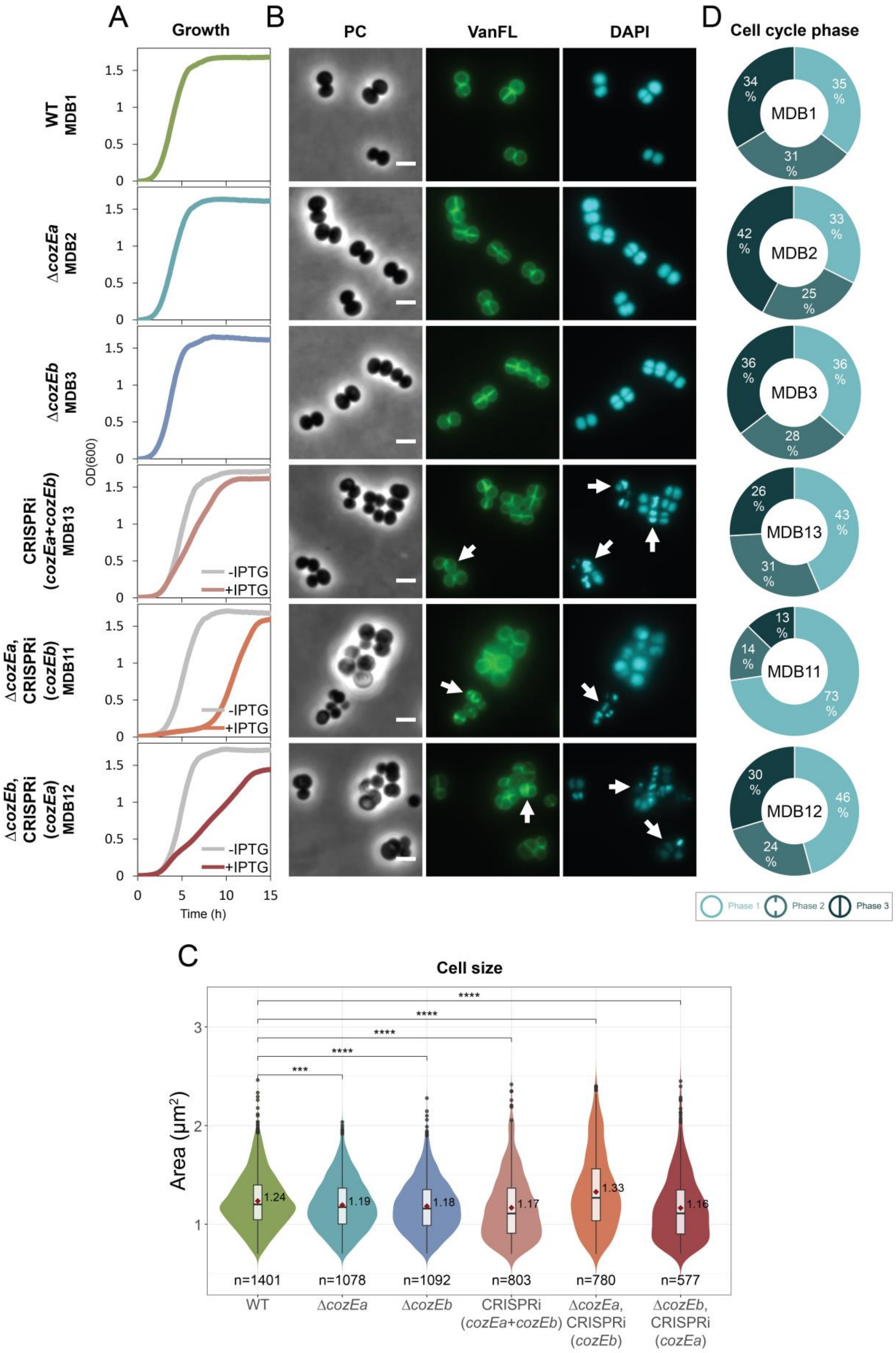
A
Growth
B
PC
VanFL
DAPI
D
Cell cycle phase
WT
MDB1
ΔcozEa
MDB2
ΔcozEb
MDB3
CRISPRi
(cozEa+cozEb)
MDB13
ΔcozEa,
CRISPRi
(cozEb)
MDB11
ΔcozEb,
CRISPRi
(cozEa)
MDB12
OD(600)
Time (h)
-IPTG
+IPTG
MDB1
MDB2
MDB3
MDB13
MDB11
MDB12
Phase 1
Phase 2
Phase 3
C
Cell size
Area (μm²)
1.24
1.19
1.18
1.17
1.33
1.16
n=1401
n=1078
n=1092
n=803
n=780
n=577
WT
ΔcozEa
ΔcozEb
CRISPRi (cozEa+cozEb)
ΔcozEa, CRISPRi (cozEb)
ΔcozEb, CRISPRi (cozEa)
Strain

**Fig. S2. Morphological and cell cycle analysis of single and double *cozE* mutants in *S. aureus* NCTC8325-4.**

(**A**) Growth curves of NCTC8325-4 wild-type (MDB1), Δ*cozEa* (MDB2) and Δ*cozEb* (MDB3), as well as of a CRISPRi double knockdown strain (CRISPRi(*cozEa*+*cozEb*), MDB13) and combined knockout/knockdown strains (Δ*cozEa*, CRISPRi(*cozEb*), MDB11, and Δ*cozEb*, CRISPRi(*cozEa*), MDB12) in BHI medium at 37°C. The graphs represent averages from triplicate measurements. The CRISPRi-strains were grown with and without IPTG, as indicated. (**B**) Micrographs of the same strains as in (A) showing phase contrast (PC) and fluorescence microscopy of cells stained with the cell wall label VanFL and the nucleoid label DAPI. CRISPRi strains were grown in medium with IPTG to induce the CRISPRi system. White arrows point to cells with perturbed septum formation and abnormal nucleoid staining. The scale bars are 2 µm. (**C**) Violin plots of the cell areas (in $\mu m^2$) of NCTC8325-4 wild-type (1.24 ± 0.27 $\mu m^2$), Δ*cozEa* (1.19 ± 0.26 $\mu m^2$), Δ*cozEb* (1.18 ± 0.27 $\mu m^2$), MDB13 (1.17 ± 0.32 $\mu m^2$), MDB11 (1.33 ± 0.39 $\mu m^2$), and MDB12 (1.16 ± 0.33 $\mu m^2$), determined using MicrobeJ. Significant differences between the strains are indicted with asterisks (* indicates a P-value of < 0.05, ** indicates a P-value of < 0.01, and *** indicates a P-value of < 0.001, derived from a Mann-Whitney test). The number of cells analyzed for each strain is indicated in the figure. (**D**) Frequency of cells in each of the three cell cycle phases for NCTC8325-4 wild-type, Δ*cozEa*, Δ*cozEb*, MDB13, MDB11, and MDB12. See Fig. 1D in the main article for a schematic overview of the different phases analyzed. The distributions were obtained by manually counting the different cell cycle phases of 100-150 randomly selected VanFL stained cells from each strain.

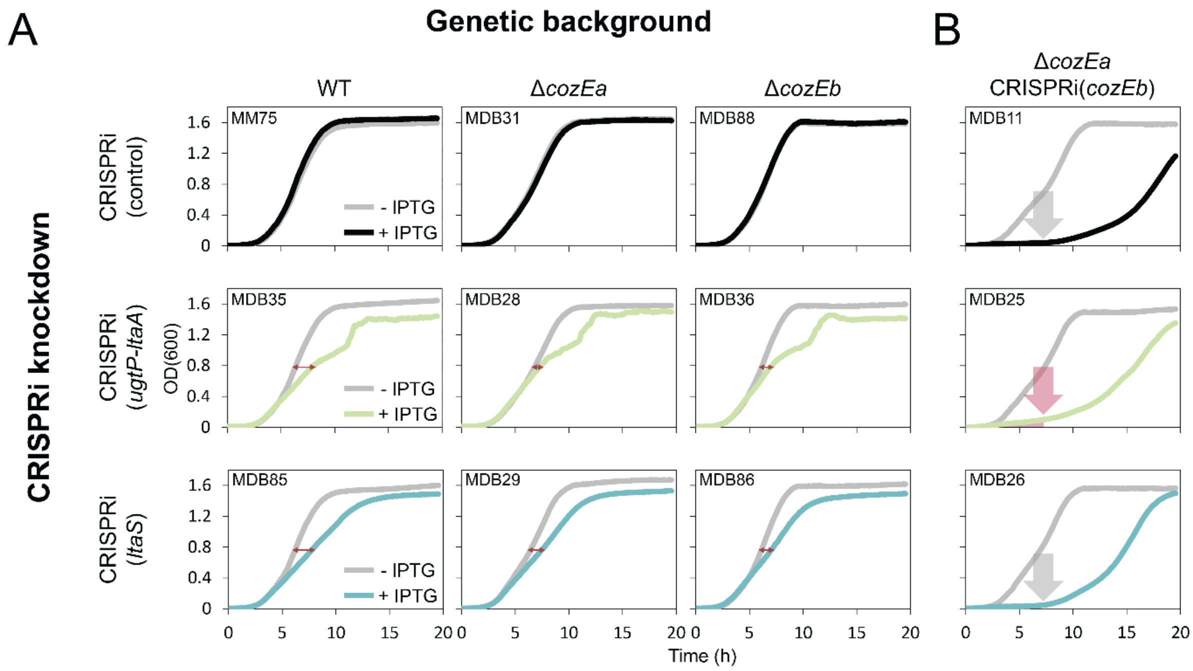


**Fig. S3**. **Synthetic genetic relationships between *cozE* genes and genes involved in LTA biosynthesis.**
(**A**) Growth assays to identify possible genetic interactions between *cozEa* or *cozEb* and the LTA biosynthetic pathway. *ugtP-ltaA* or *ltaS* was knocked down by CRISPRi in either a NCTC8325-4 wild-type, Δ*cozEa*, or Δ*cozEb* background. The CRISPRi-strains were grown with and without IPTG, as indicated. The length of the red arrows illustrates the growth differences observed between the genetic backgrounds, wild-type, Δ*cozEa*, and Δ*cozEb*, when UgtP-LtaA or LtaS were depleted. (**B**) Growth assays to identify possible genetic interactions between *cozEab* and the LTA biosynthetic pathway. *ugtP-ltaA* or *ltaS* was knocked down along with *cozEb* in a Δ*cozEa* background. The CRISPRi-strains were grown with and without IPTG, as indicated by the colors. The red arrow points to the slight growth alleviation observed when *ugtP-ltaA* was knocked down together with *cozEb* in the Δ*cozEa* background (MDB25). In both (A) and (B), the graphs represent averages from triplicate measurements.

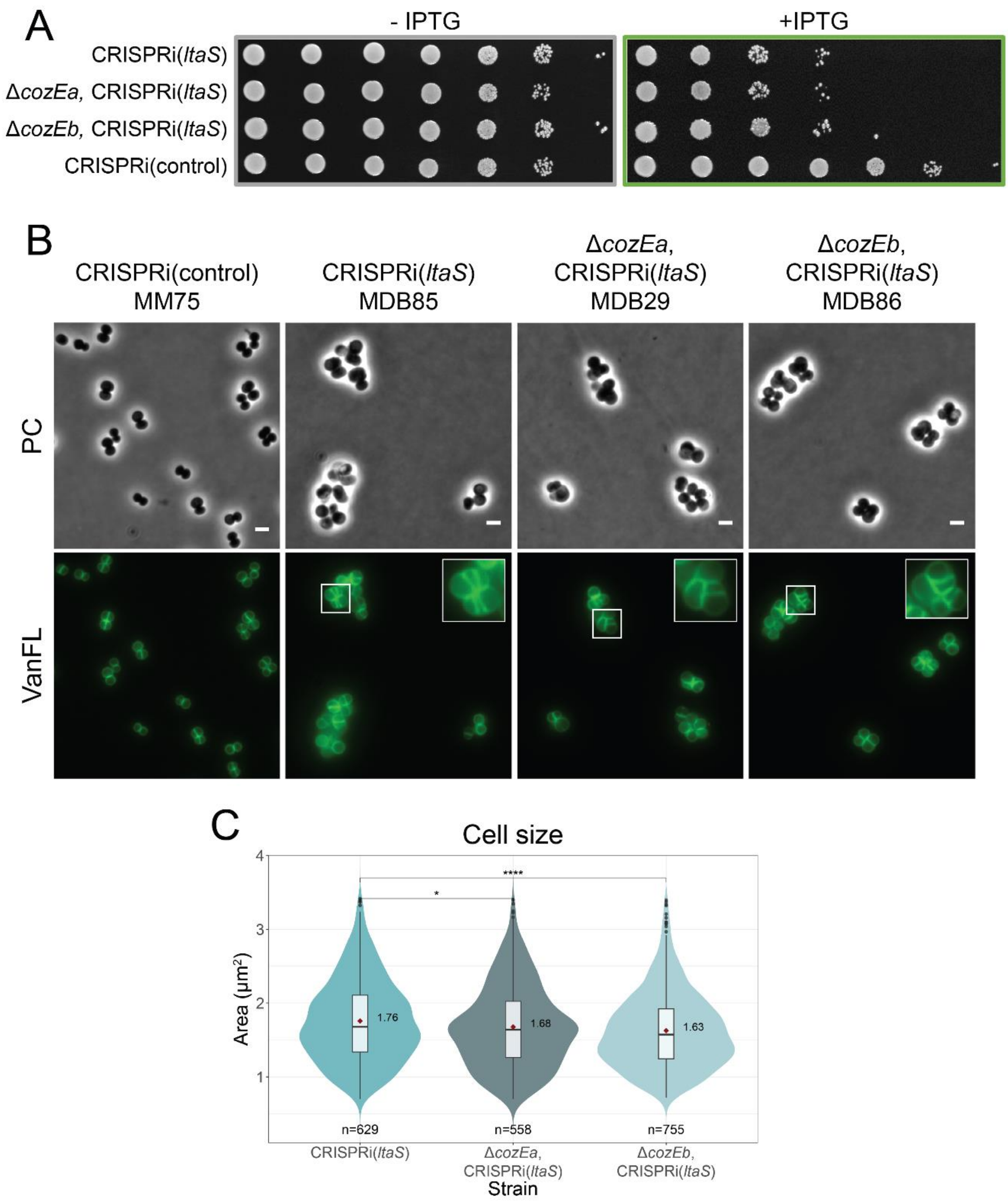


**Fig. S4. LtaS depletion in wild-type, Δ*cozEa*, and Δ*cozEb* *S. aureus* NCTC8325-4 cells.**
(**A**) Growth on solid medium of LtaS depleted *S. aureus* NCTC8325-4 cells; CRISPRi(*ltaS*) (MDB85), Δ*cozEa*, CRISPRi(*ltaS*) (MDB29), and Δ*cozEb*, CRISPRi(*ltaS*) (MDB86). In addition to a strain carrying a non-targeting sgRNA (MM75) used as a control. 10-fold dilution series, made from noninduced overnight cultures, were spotted onto agar plates with and without IPTG, as indicated. (**B**) Phase contrast (PC) and VanFL staining micrographs of the same strains as in (A). The cells were grown in the presence of IPTG for induction of the CRISPRi system. The scale bars are 2 µm. (**C**) Violin plots of the cell areas (in $\mu m^2$) of NCTC8325-4 CRISPRi(*ltaS*) (1.76 ± 0.55 $\mu m^2$), Δ*cozEa*, CRISPRi(*ltaS*) (1.68 ± 0.55), and Δ*cozEb*, CRISPRi(*ltaS*) (1.63 ± 0.51 $\mu m^2$), determined using MicrobeJ. Significant differences between the strains are indicted with asterisks (* indicates a P-value of < 0.05, ** indicates a P-value of < 0.01, and *** indicates a P-value of < 0.001, derived from a Mann-Whitney test). The number of cells analyzed for each strain is indicated in the figure.

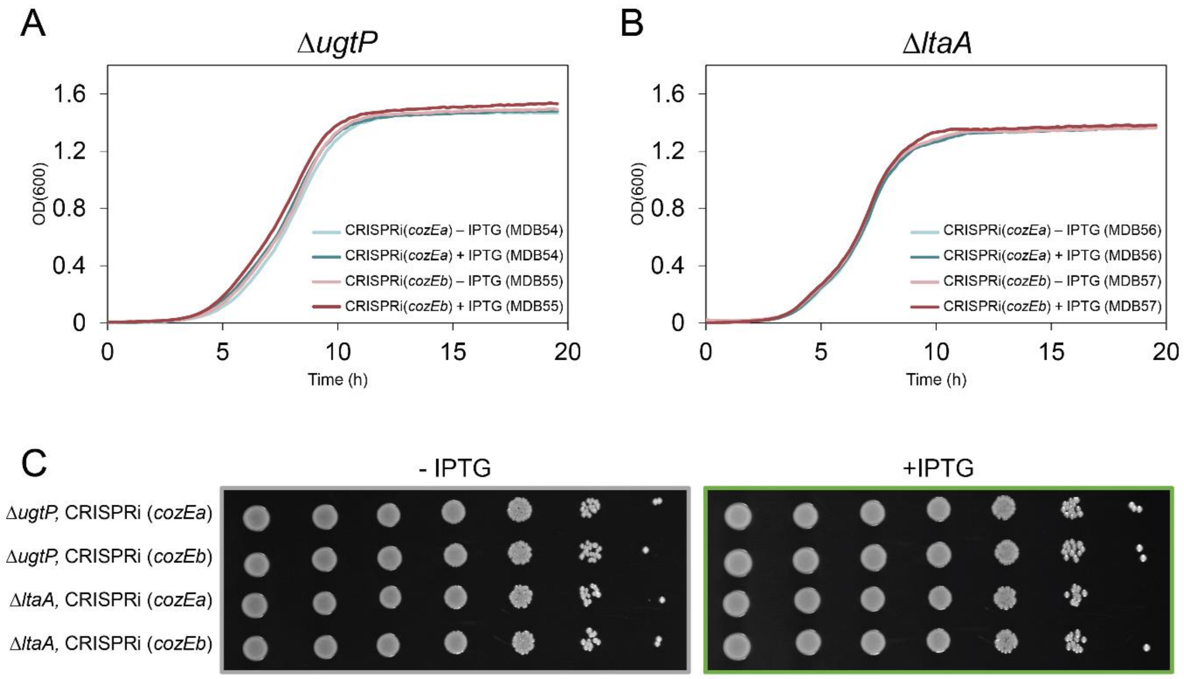


**Fig. S5. Single CozE depletion in *S. aureus* JE2 Δ*ugtP* and Δ*ltaA*.**

Growth curves of JE2 (**A**) Δ*ugtP* and (**B**) Δ*ltaA* with individual depletion of CozEa or CozEb in BHI medium at 37°C. The graphs represent averages from triplicate measurements. The CRISPRi-strains were grown with and without IPTG, as indicated. (**C**) Additionally, growth of the JE2 Δ*ugtP* and Δ*ltaA* strains with single CozE depletion on solid medium. 10-fold dilution series, made from noninduced overnight cultures, were spotted onto agar plates with and without IPTG, as indicated.

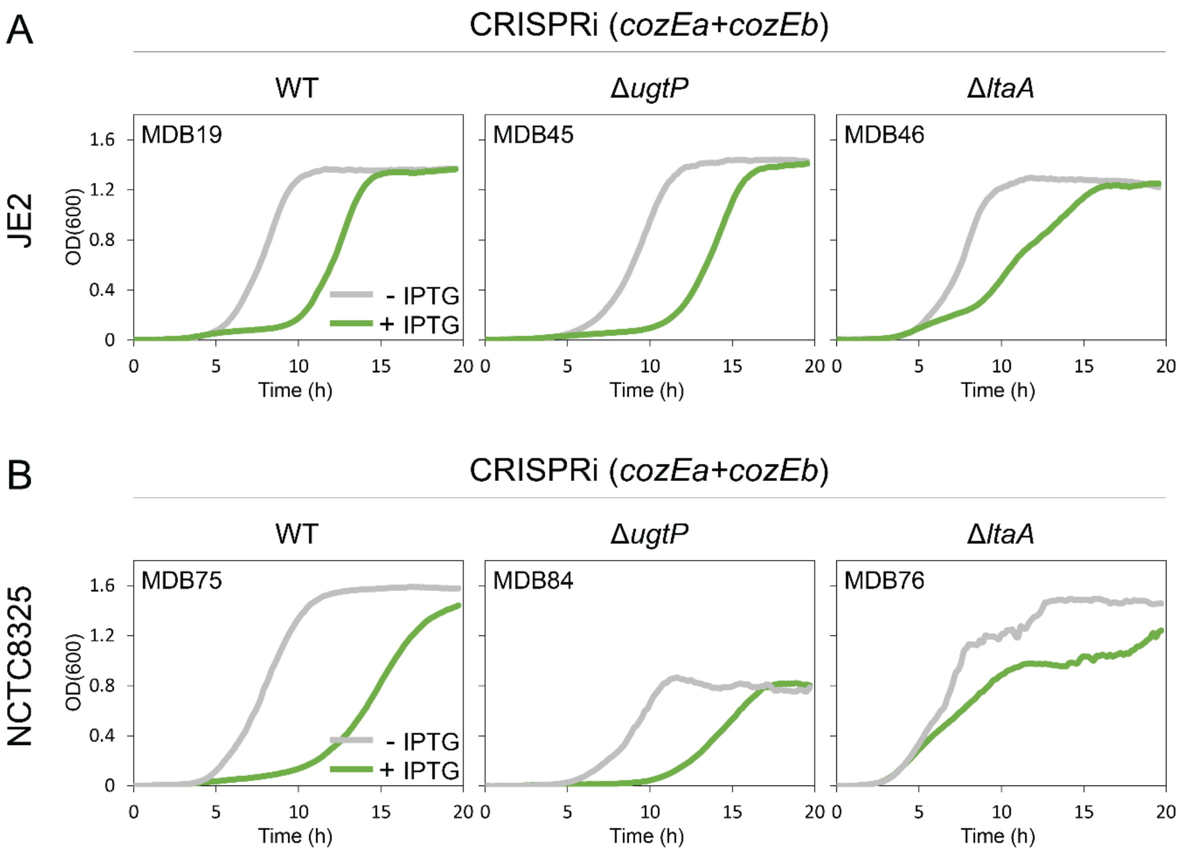


**Fig. S6**. **Growth of wild-type, Δ*ugtP*, and Δ*ltaA* with double CozE depletion in *S. aureus* JE2 and NCTC8325, using 500 µM IPTG for maximum depletion.**

Growth of wild-type, Δ*ugtP*, and Δ*ltaA* with double *cozE* knockdown in (**A**) JE2 (MDB19, MDB45, and MDB46) and (**B**) NCTC8325 (MDB75, MDB84, MDB76) in liquid cultures. Cells were grown in the presence or absence of 500 µM IPTG, as indicated by the colors. The graphs represent averages from triplicate measurements.

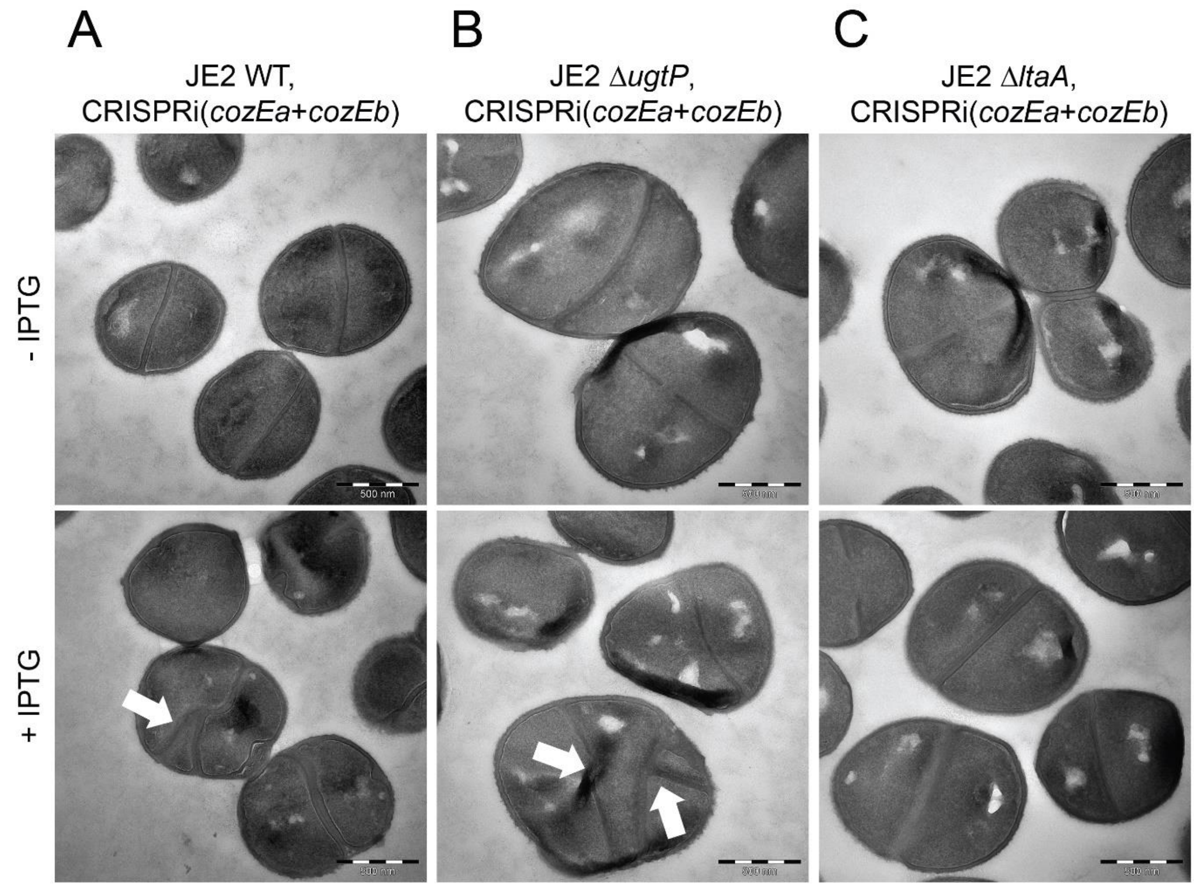


**Fig. S7. TEM analysis of *S. aureus* JE2 wild-type, Δ*ugtP*, and Δ*ltaA* mutants with double CozE depletion.** TEM micrographs of JE2 (**A**) wild-type, (**B**) Δ*ugtP*, and (**C**) Δ*ltaA* cells with uninduced or induced depletion of CozEa and CozEb. White arrows point to cells with aberrant septum formation. The scale bars are 500 nm.

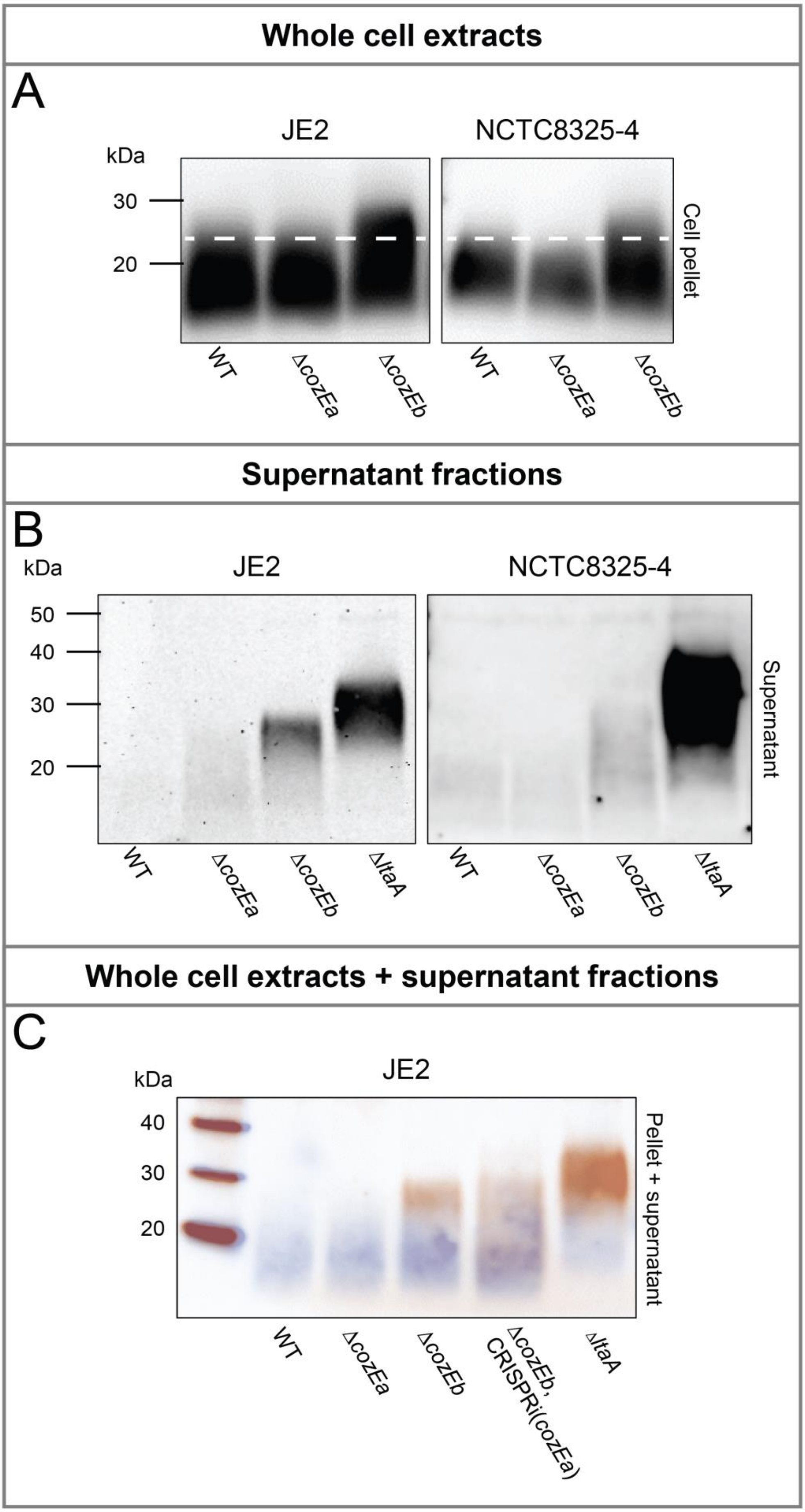
Whole cell extracts
A
JE2
NCTC8325-4
kDa
30
20
Cell pellet
WT
ΔcozEa
ΔcozEb
WT
ΔcozEa
ΔcozEb
Supernatant fractions
B
kDa
JE2
NCTC8325-4
50
40
30
20
Supernatant
WT
ΔcozEa
ΔcozEb
ΔltaA
WT
ΔcozEa
ΔcozEb
ΔltaA
Whole cell extracts + supernatant fractions
C
JE2
kDa
40
30
20
Pellet + supernatant
WT
ΔcozEa
ΔcozEb
ΔcozEb, CRISPRi(cozEa)
ΔltaA

**Fig. S8. Characterization of LTA polymer length and stability in single and double *cozE* mutants.**
LTA polymers were detected in whole cell extracts, (A) and (C), or in supernatant fractions, (B) and (C), by immunoblotting with an anti-LTA antibody. (**A**) Immunoblots of wild-type, Δ*cozEa*, and Δ*cozEb* in JE2 (MDB37, MDB38, and MDB10) and NCTC8325-4 (MDB1, MDB2, and MDB3). (**B**) Immunoblots of wild-type, Δ*cozEa*, Δ*cozEb*, and Δ*ltaA* in JE2 (MDB9, MDB38, MDB10, and MDB40) and NCTC8325-4 (MDB1, MDB2, MDB3, and MDB69). (**C**) Merged immunoblots of JE2 wild-type (MDB9), Δ*cozEa* (MDB38), Δ*cozEb* (MDB10), a double *cozE* mutant (MDB21), and a positive control strain (MDB40), where the LTA detected in the cell pellets are colored blue while the LTA detected in the supernatants are colored orange to illustrate the differences in LTA polymer lengths.

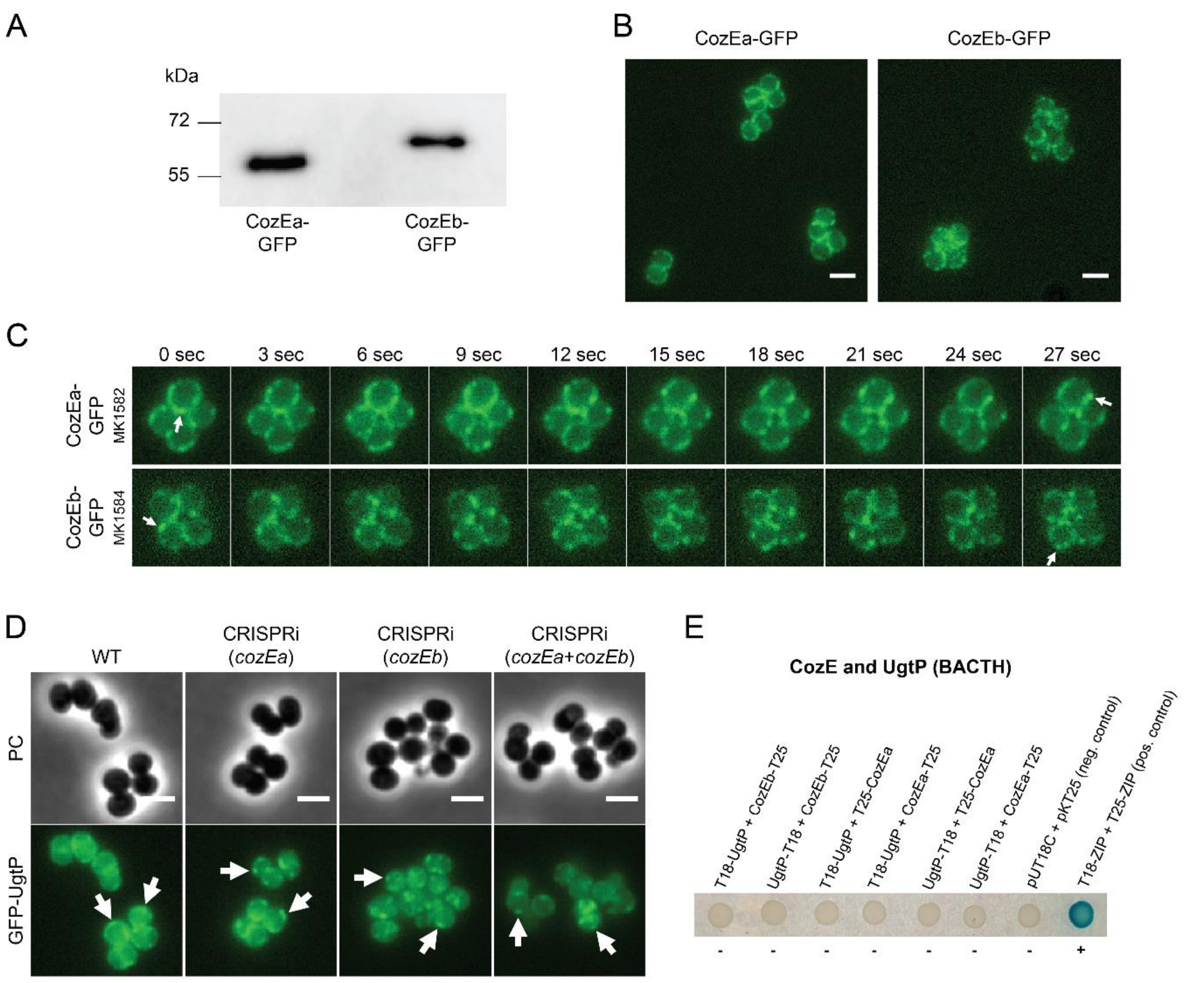


**Fig. S9. Subcellular localization of CozEa, CozEb and UgtP.**

(**A**) The relative expression of CozEa and CozEb is indicated by the band density of CozEa-GFP and CozEb-GFP (from MK1582 and MK1584, respectively) in an immunoblot assay using an anti-GFP antibody. (**B**) The subcellular localization of CozEa and CozEb analyzed by florescent microscopy of MK1582 and MK1584, respectively. The scale bars are 2 µm. (**C**) The movement of the CozE proteins were analyzed by time-lapse florescent microscopy of MK1582 (with a *cozEa-gfp* fusion) and MK1584 (with a *cozEb-gfp* fusion). Images were captured every third second (x10), as indicated in the figure. White arrows highlight the membrane movement of CozEa-GFP and CozEb-GFP, as the signals pointed to in the initial images are no longer present at the same location in the membrane after 27 seconds. The dynamic spatiotemporal localization of CozEa and CozEb is further depicted in **Movie S1** and **S2**, respectively. (**D**) Localization of GFP-UgtP in NCTC8325-4 wild-type (MDB77), as well as cells depleted of CozEa (MDB89), CozEb (MDB90), or both CozE proteins (MDB79). Phase contrast- and fluorescence images are shown. Arrows point to spots with membrane-localized GFP-UgtP. The scale bars are 2 µm. (**E**) Bacterial two-hybrid interaction assays between CozEa or CozEb and UgtP. Fusions of the proteins to either the T18 or K25 domain of adenylate cyclase were expressed in *E. coli* according to established protocols (1). Spots with a blue color, marked with a plus sign, indicate positive interactions, while white spots, marked with a minus symbol, indicate no interaction. Positive and negative controls are included. The presented BATCH assays are representative of 5 independent replicates.

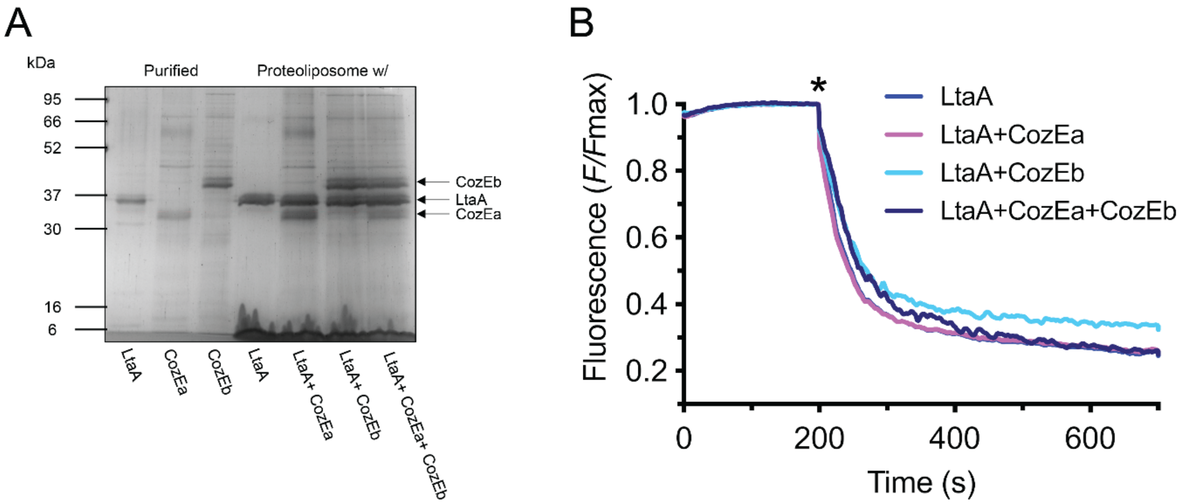


**Fig. S10. LtaA-catalyzed Glc$_2$DAG flipping in the presence of CozE proteins.**
(**A**) SDS-PAGE of purified (1) LtaA (44.6 kDa), (2) CozEa (40.1 kDa), and (3) CozEb (45.1 kDa), in addition to proteoliposomes with (1) LtaA, (2) LtaA and CozEa, (3) LtaA and CozEb, and (4) LtaA, CozEa, and CozEb used in the Glc$_2$DAG flipping experiment. Arrows point at LtaA, CozEa, and CozEb. (**B**) Representative traces of proteoliposomes containing LtaA and LtaA together with CozE proteins (n = 3). The asterisk marks addition of dithionite. F corresponds to the fluorescence intensity measured for each time point. Fmax is the average fluorescence measured during the first 200 seconds of the experiment.

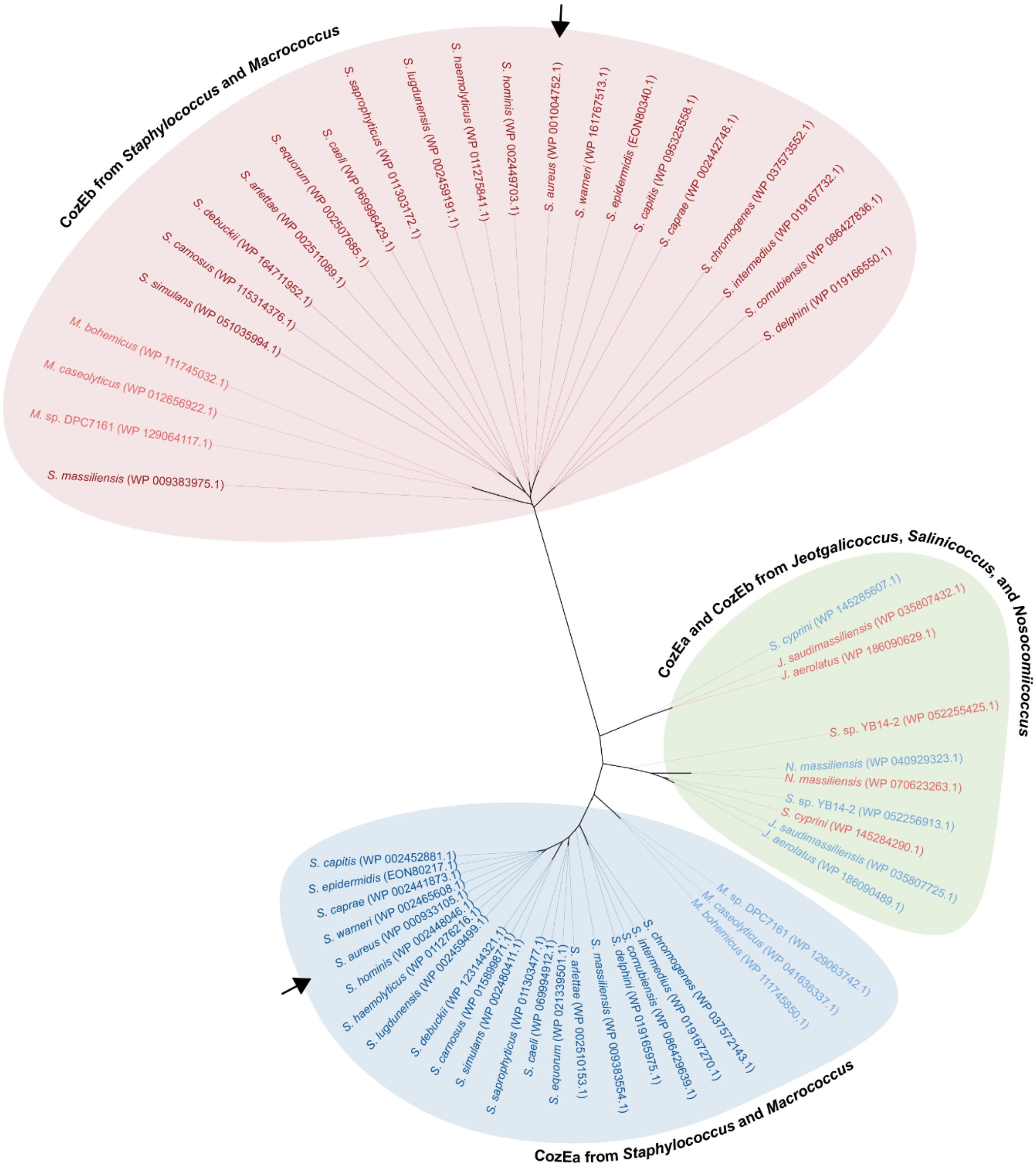


**Fig. S11. Phylogenetic distribution of CozE proteins in the *Staphylococcaceae* family.**

A maximum likelihood phylogenetic tree was constructed from a Clustal Omega sequence alignment of 56 CozE homologs from the *Staphylococcaceae* family, specifically *Staphylococcus*, *Macrococcus*, *Jeotgalicoccus*, *Salinicoccus*, and *Nosocomiicoccus*, using IQ-TREE. The phylogenetic tree was visualized and annotated using iTOL. The CozE proteins from species belonging to *Staphylococcus* and *Macrococcus* distributed into two phylogenetically separate subgroups; CozEa (marked in blue) and CozEb (marked in red). The CozE proteins from *Jeotgalicoccus*, *Salinicoccus*, and *Nosocomiicoccus* clustered together in another subgroup (marked in green), which is phylogenetically closer to the CozEa subgroup than the CozEb subgroup. Arrows point at the CozE proteins found in *S. aureus*.

### Supplementary movies

**Movie S1 (separate file). The dynamic spatiotemporal localization of GFP-tagged CozEa.**
The movement of CozEa was analyzed by time-lapse fluorescent microscopy of MK1582, a NCTC8325-4 mutant with a *cozEa-gfp* fusion gene chromosomally integrated in its native locus. Images were captured every third second (x10), as indicated in the movie.

**Movie S2 (separate file). The dynamic spatiotemporal localization of GFP-tagged CozEb.**
The movement of CozEb was analyzed by time-lapse fluorescent microscopy of MK1584, a NCTC8325-4 mutant with a *cozEb-gfp* fusion gene chromosomally integrated in its native locus. Images were captured every third second (x10), as indicated in the movie.

### Supplementary tables

**Table S1.** Strains and mutants used in this work.

| Name | Genotype and characteristics[a] | Reference |
|---|---|---|
| ***S. aureus* JE2** | | |
| JE2/MDB9/ MDB37 | Community acquired MRSA strain, derivative of USA300 LAC cured of plasmids | (2) |
| MDB38/ NE1270 | JE2 Δ*cozEa*, ery$^r$ | (2) |
| MDB10/ NE779 | JE2 Δ*cozEb*, ery$^r$ | (2) |
| MDB39/ NE1663 | JE2 Δ*ugtP*, ery$^r$ | (2) |
| MDB40/ NE462 | JE2 Δ*ltaA*, ery$^r$ | (2) |
| MDB16 | JE2 carrying pLOW-dCas9_aad9, spc$^r$ | This work |
| MDB20 | MDB10 carrying pLOW-dCas9_aad9, spc$^r$ | This work |
| MDB41 | MDB39 carrying pLOW-dCas9_aad9, spc$^r$ | This work |
| MDB42 | MDB40 carrying pLOW-dCas9_aad9, spc$^r$ | This work |
| MDB17 | MDB16 carrying pCG248-sgRNA(*cozEa*), spc$^r$, cam$^r$ | This work |
| MDB18 | MDB16 carrying pCG248-sgRNA(*cozEb*), spc$^r$, cam$^r$ | This work |
| MDB19 | MDB16 carrying pCG248-sgRNA(*cozEa*+*cozEb*), spc$^r$, cam$^r$ | This work |
| MDB44 | MDB16 carrying pCG248-sgRNA(*luc*), spc$^r$, cam$^r$ | This work |
| MDB21 | MDB20 carrying pCG248-sgRNA(*cozEa*+*cozEb*), spc$^r$, cam$^r$ | This work |
| MDB45 | MDB41 carrying pCG248-sgRNA(*cozEa*+*cozEb*), spc$^r$, cam$^r$ | This work |
| MDB46 | MDB42 carrying pCG248-sgRNA(*cozEa*+*cozEb*), spc$^r$, cam$^r$ | This work |
| MDB54 | MDB41 carrying pCG248-sgRNA(*cozEa*), spc$^r$, cam$^r$ | This work |
| MDB55 | MDB41 carrying pCG248-sgRNA(*cozEb*), spc$^r$, cam$^r$ | This work |
| MDB56 | MDB42 carrying pCG248-sgRNA(*cozEa*), spc$^r$, cam$^r$ | This work |
| MDB57 | MDB42 carrying pCG248-sgRNA(*cozEb*), spc$^r$, cam$^r$ | This work |
| MDB59 | MDB10 carrying pRAB11-*cozEa*, cam$^r$ | This work |
| MDB60 | MDB10 carrying pRAB11-*cozEb*, cam$^r$ | This work |
| MDB176 | MDB10 carrying pRAB11, cam$^r$ | This work |
| ***S. aureus* NCTC8325-4** | | |
| NCTC8325-4/ MDB1 | MSSA lab strain, derivative of NCTC8325 cured of prophages | (3) |
| MDB2 | NCTC8325-4 Δ*cozEa*, spc$^r$ | This work |
| MDB3 | NCTC8325-4 Δ*cozEb*, spc$^r$ | This work |
| MH225 | NCTC8325-4 carrying pLOW-dCas9_extra_*lacO*, ery$^r$ | Lab collection |
| MH223 | MDB2 carrying pLOW-dCas9_extra_*lacO*, ery$^r$ | This work |
| MH224 | MDB3 carrying pLOW-dCas9_extra_*lacO*, ery$^r$ | This work |
| MDB11 | MH223 carrying pCG248-sgRNA(*cozEb*), ery$^r$, cam$^r$ | This work |
| MDB12 | MH224 carrying pCG248-sgRNA(*cozEa*), ery$^r$, cam$^r$ | This work |
| MDB13 | MH225 carrying pCG248-sgRNA(*cozEa*+*cozEb*), ery$^r$, cam$^r$ | This work |
| MDB14 | MH225 carrying pCG248-sgRNA(*cozEa*), ery$^r$, cam$^r$ | This work |
| MDB15 | MH225 carrying pCG248-sgRNA(*cozEb*), ery$^r$, cam$^r$ | This work |
| MM75 | MH225 carrying pVL2336-sgRNA(*luc*), ery$^r$, cam$^r$ | Lab collection |
| MDB31 | MH223 carrying pCG248-sgRNA(*luc*), ery$^r$, cam$^r$ | This work |
| MDB88 | MH224 carrying pCG248-sgRNA(*luc*), ery$^r$, cam$^r$ | This work |

| | | |
|---|---|---|
| MDB25 | MH223 carrying pCG248-sgRNA(*cozEb+ugtP-ltaA*), $ery^r$, $cam^r$ | This work |
| MDB26 | MH223 carrying pCG248-sgRNA(*cozEb+ltaS*), $ery^r$, $cam^r$ | This work |
| MDB35 | MH225 carrying pVL2336-sgRNA(*ugtP-ltaA*), $ery^r$, $cam^r$ | This work |
| MDB28 | MH223 carrying pVL2336-sgRNA(*ugtP-ltaA*), $ery^r$, $cam^r$ | This work |
| MDB36 | MH224 carrying pVL2336-sgRNA(*ugtP-ltaA*), $ery^r$, $cam^r$ | This work |
| MDB85 | MH225 carrying pVL2336-sgRNA(*ltaS*), $ery^r$, $cam^r$ | This work |
| MDB29 | MH223 carrying pVL2336-sgRNA(*ltaS*), $ery^r$, $cam^r$ | This work |
| MDB86 | MH224 carrying pVL2336-sgRNA(*ltaS*), $ery^r$, $cam^r$ | This work |
| MDB58 | MDB3 carrying pRAB11-*cozEa*, $cam^r$ | This work |
| MDB62 | MDB3 carrying pRAB11-*cozEb*, $cam^r$ | This work |
| MDB174 | MDB3 carrying pRAB11, $cam^r$ | This work |
| MK1582 | NCTC8325-4, but with *gfp* fused to the 3' end of *cozEa*, $spc^r$ | This work |
| MK1584 | NCTC8325-4, but with *gfp* fused to the 3' end of *cozEb*, $spc^r$ | This work |
| MDB77 | NCTC8325-4, but with *gfp* fused to the 5' end of *ugtP*, $spc^r$ | This work |
| MDB78 | MDB77 carrying pLOW-dCas9_extra_*lacO*, $ery^r$ | This work |
| MDB79 | MDB78 carrying pCG248-sgRNA(*cozEa+cozEb*), $ery^r$, $cam^r$ | This work |
| MDB89 | MDB78 carrying pCG248-sgRNA(*cozEa*), $ery^r$, $cam^r$ | This work |
| MDB90 | MDB78 carrying pCG248-sgRNA(*cozEb*), $ery^r$, $cam^r$ | This work |
| ***S. aureus* NCTC8325** | | |
| NCTC8325/ MDB68 | MSSA lab strain | Lab collection |
| MDB80 | NCTC8325 Δ*ugtP*, $spc^r$ | This work |
| MDB69/ VL3222 | NCTC8325 Δ*ltaA*, $spc^r$ | (4) |
| MDB70 | NCTC8325 carrying pLOW-dCas9_extra_*lacO*, $ery^r$ | This work |
| MDB81 | MDB80 carrying pLOW-dCas9_extra_*lacO*, $ery^r$ | This work |
| MDB71 | MDB69 carrying pLOW-dCas9_extra_*lacO*, $ery^r$ | This work |
| MDB75 | MDB70 carrying pCG248-sgRNA(*cozEa+cozEb*), $ery^r$, $cam^r$ | This work |
| MDB84 | MDB81 carrying pCG248-sgRNA(*cozEa+cozEb*), $ery^r$, $cam^r$ | This work |
| MDB76 | MDB71 carrying pCG248-sgRNA(*cozEa+cozEb*), $ery^r$, $cam^r$ | This work |
| ***S. aureus* SH1000** | | |
| SH1000 | MSSA lab strain, *rsbU*$^+$ *agr*$^+$ derivative of NCTC8325-4 | (5) |
| ***E. coli*** | | |
| IM08B | DH10B, Δ*dcm*, $P_{help}$-*hsdMS*, $P_{N25}$-*hsdS* (strain expressing the *S. aureus* CC8 specific methylation genes) | (6) |
| XL1-Blue | K-12 $lacI^q$ strain used for BACTH plasmid preparation | Agilent |
| BTH101 | Δ*cya* strain used for BACTH analysis | Euromedex |
| BL21 Gold (DE3) | $F^-$ *ompT hsdS*($r_B^-$ $m_B^-$) *dcm*$^+$ $Tet^r$ *gal* λ(DE3) *endA* Hte used for expression of LtaA and CozE for the *in vitro* flipping assay | Stratagene |

a. $ery^r$ = erythromycin resistant, $spc^r$ = spectinomycin resistant, and $cam^r$ = chloramphenicol resistant.

**Table S2.** Plasmids used in this work.

| Name | Description[a] | Reference |
|---|---|---|
| pLOW | Low-copy number staphylococcal shuttle vector with a IPTG inducible P*spac* promoter and *lacI* repressor, amp$^{r}$, ery$^{r}$ | (7) |
| pLOW-*dCas9_aad9* | For IPTG inducible expression of dCas9, amp$^{r}$, spc$^{r}$ | (8) |
| pLOW-*dCas9*_extra_*lacO* | For IPTG inducible expression of dCas9, amp$^{r}$, ery$^{r}$ | (9) |
| pLOW-*cozEa-m(sf)gfp* | For IPTG inducible expression of CozEa with GFP fused to its C-terminal, amp$^{r}$, ery$^{r}$ | (9) |
| pLOW-*cozEb-m(sf)gfp* | For IPTG inducible expression of CozEb with GFP fused to its C-terminal, amp$^{r}$, ery$^{r}$ | (9) |
| pLOW-*m(sf)gfp*-SA1477 | For IPTG inducible expression of SAOUHSC_1477 with GFP fused to its N-terminal, amp$^{r}$, ery$^{r}$ | Lab collection |
| pLOW- *m(sf)gfp-ugtP* | For IPTG inducible expression of UgtP with GFP fused to its N-terminal, amp$^{r}$, ery$^{r}$ | This work |
| pCG248 | *E. coli*/*S. aureus* shuttle vector, amp$^{r}$, cam$^{r}$ | (10) |
| pCG248-sgRNA(*cozEa*) | For constitutive expression of sgRNA(*cozEa*), amp$^{r}$, cam$^{r}$ | (9) |
| pCG248-sgRNA(*cozEb*) | For constitutive expression of sgRNA(*cozEb*), amp$^{r}$, cam$^{r}$ | (9) |
| pCG248-sgRNA(*cozEa*+*cozEb*) | For constitutive expression of sgRNA(*cozEa*+*cozEb*), amp$^{r}$, cam$^{r}$ | (9) |
| pCG248-sgRNA(*cozEb*+*ugtP-ltaA*) | For constitutive expression of sgRNA(*cozEb*+*ugtP-ltaA*), amp$^{r}$, cam$^{r}$ | This work |
| pCG248-sgRNA(*cozEb*+*ltaS*) | For constitutive expression of sgRNA(*cozEb*+*ltaS*), amp$^{r}$, cam$^{r}$ | This work |
| pCG248-sgRNA(*luc*) | For constitutive expression of sgRNA(control), amp$^{r}$, cam$^{r}$ | (9) |
| pVL2336 | *E. coli*/*S. aureus* shuttle vector, amp$^{r}$, cam$^{r}$ | (11) |
| pVL2336-sgRNA(*ugtP-ltaA*) | For constitutive expression of sgRNA(*ugtP-ltaA*), amp$^{r}$, cam$^{r}$ | This work |
| pVL2336-sgRNA(*ltaS*) | For constitutive expression of sgRNA(*ltaS*), amp$^{r}$, cam$^{r}$ | This work |
| pMAD | Thermosensitive shuttle vector for allelic replacement in Gram-positive bacteria, amp$^{r}$, ery$^{r}$ | (12) |
| pMAD-*cozEa*::*spc* | For allelic replacement of *cozEa*, amp$^{r}$, ery$^{r}$, spc$^{r}$ | (9) |
| pMAD-*cozEb*::*spc* | For allelic replacement of *cozEb*, amp$^{r}$, ery$^{r}$, spc$^{r}$ | (9) |
| pMAD-*cozEa*::*cam* | For allelic replacement of *cozEa*, amp$^{r}$, ery$^{r}$, cam$^{r}$ | (9) |
| pMAD-Δ*ugtP*::*spc* | For allelic replacement of *ugtP*, amp$^{r}$, ery$^{r}$, spc$^{r}$ | This work |
| pMAD-*cozEa-m(sf)gfp_spc* | To GFP-tag *cozEa* in its native locus, amp$^{r}$, ery$^{r}$, spc$^{r}$ | This work |
| pMAD-*cozEb-m(sf)gfp_spc* | To GFP-tag *cozEb* in its native locus, amp$^{r}$, ery$^{r}$, spc$^{r}$ | This work |
| pMAD-P$_{ugtP}$-*m(sf)gfp-ugtP_spc* | To GFP-tag *ugtP* in a natural locus under the control of its native promotor, amp$^{r}$, ery$^{r}$, spc$^{r}$ | This work |
| pRAB11 | *E. coli*/*S. aureus* shuttle vector with an anhydrotetracycline inducible *xyl*/*tet* promoter, amp$^{r}$, cam$^{r}$ | (10) |
| pRAB11-*cozEa* | For aTc inducible expression of CozEa, amp$^{r}$, cam$^{r}$ | This work |
| pRAB11-*cozEb* | For aTc inducible expression of CozEa, amp$^{r}$, cam$^{r}$ | This work |
| pKT25 | Vector encoding the T25 fragment (residues 1-224) of adenylate cyclase, CyaA, upstream of a MCS, under the transcriptional control of a *lac* promoter, kan$^{r}$ | Euromedex |
| pKT25-*cozEa* | For expression of CozEa fused in frame to the C-terminal end of T25, kan$^{r}$ | (9) |
| pKT25-*cozEb* | For expression of CozEb fused in frame to the C-terminal end of T25, kan$^{r}$ | (9) |

| | | |
|---|---|---|
| pKT25-*zip* | For expression of the leucine zipper of GCN4 fused in frame to the C-terminal end of T25, kan$^r$ | Euromedex |
| pKNT25 | Vector encoding the T25 fragment (residues 1-224) of adenylate cyclase, CyaA, downstream of a MCS, under the transcriptional control of a *lac* promoter, kan$^r$ | Euromedex |
| pKNT25-*cozEa* | For expression of CozEa fused in frame to the N-terminal end of T25, kan$^r$ | (9) |
| pKNT25-*cozEb* | For expression of CozEb fused in frame to the N-terminal end of T25, kan$^r$ | (9) |
| pUT18 | Vector encoding the T18 fragment (residues 225-399) of adenylate cyclase, CyaA, downstream of a MCS, under the transcriptional control of a *lac* promoter, amp$^r$ | Euromedex |
| pUT18-*cozEa* | For expression of CozEa fused in frame to the N-terminal end of T18, amp$^r$ | (9) |
| pUT18-*ugtP* | For expression of UgtP fused in frame to the N-terminal end of T18, amp$^r$ | This work |
| pUT18C | Vector encoding the T18 fragment (residues 225-399) of adenylate cyclase, CyaA, upstream of a MCS, under the transcriptional control of a *lac* promoter, amp$^r$ | Euromedex |
| pUT18C-*cozEa* | For expression of CozEa fused in frame to the C-terminal end of T18, amp$^r$ | (9) |
| pUT18C-*cozEb* | For expression of CozEb fused in frame to the C-terminal end of T18, amp$^r$ | (9) |
| pUT18C-*ugtP* | For expression of UgtP fused in frame to the C-terminal end of T18, amp$^r$ | This work |
| pUT18C-*zip* | For expression of the leucine zipper of GCN4 fused in frame to the C-terminal end of T18, amp$^r$ | Euromedex |
| pCN55 | *E. coli*/*S. aureus* shuttle vector, amp$^r$, spc$^r$ | (13) |
| LtaA-pET19b | For expression and purification of LtaA, amp$^r$ | (14) |
| pET19b-*cozEa* | For expression and purification of CozEa, amp$^r$ | This study |
| pET19b-*cozEb* | For expression and purification of CozEb, amp$^r$ | This study |

a. amp$^r$ = ampicillin resistant, ery$^r$ = erythromycin resistant, spc$^r$ = spectinomycin resistant, cam$^r$ = chloramphenicol resistant, and kan$^r$ = kanamycin resistant.

**Table S3.** Primers used in this work.

| Name | Sequence 5'-3'[a] | Description[b] |
|---|---|---|
| **Primers to check for the presence of *cozEa*** | | |
| im17 | ATCGGTACCCAATAAAACTAGGAGGAAATTTAAATGTTAAACAAGGTTTGGTTCC | *cozEa* F w/ KpnI RS |
| im18 | GATGAATTCTTAGTCCTTAACATTACTGTTTG | *cozEa* R w/ EcoRI RS |
| **Primers to check for the deletion of *cozEa*** | | |
| mk188 | ATTGGGCCCACCTAGGATC | F upstream of *cozEa* deletion |
| mk187 | CAAACATTTATCGTTGTAATACGT | R downstream of *cozEa* deletion |
| **Primers to check for the presence of *cozEb*** | | |
| gs653 | GATCGGATCCCAATGAAAATGAAAAGAATATAAGAAAG | *cozEb* F w/BamHI RS |
| gs654 | GATCGAATCCTTTATTCAACTATTTTATTACTTTCTTTA | *cozEb* R |
| **Primers to check for the deletion of *cozEb*** | | |
| mk188 | ATTGGGCCCACCTAGGATC | F upstream of *cozEb* deletion |
| mk195 | GCGTCAACAATTACACCACAG | R downstream of *cozEb* deletion |
| **Primers check for the presence of pLOW plasmids** | | |
| im218 | TCTCATTCAATTCCTAGGTGG | pLOW F |
| im134 | TGTGCTGCAAGGCGATTAAG | pLOW R |
| **Primers to check for the presence of pCG248/pVL2336 plasmids** | | |
| mk26 | GGATAACCGTATTACCGCCT | pCG248 F |
| mk25 | AAATCTCGAAAATAATAGAGGGA | pCG248 R |
| **Primers to check for the presence of pRAB11 plasmids** | | |
| mk23 | GGATCCCCTCGAGTTCATG | pRAB11 F |
| mk24 | GGGATGTGCTGCAAGGCGA | pRAB11 R |
| **Primers to check for the presence of pMAD plasmids** | | |
| im156 | AATCTAGCTAATGTTACGTTACA | pMAD F |
| mk177 | GATGCCGCCGGAAGCGAG | pMAD R |
| **Primers for construction of pLOW-*m(sf)gfp-ugtP*** | | |
| mdb9 | ACGTGGATCCGTTACTCAAAATAAAAAGATATTGA | *ugtP* F w/ BamHI RS |
| mdb2 | ACGTGAATTCATGATTAGCGTAATTATTTAACG | *ugtP* R w/ EcoRI RS |
| **Primers for construction of pMAD-$P_{ugtP}$-*m(sf)gfp-ugtP_spc*** | | |
| mdb3 | ACCTGAATTCGGTATCGCTAGCGATGGCT | ori_up F w/ EcoRI RS |
| mdb4 | TCGAACCCCGATGTTGTCG | ori_up R |
| mdb5 | **CGACAACATCGGGGTTCGA**CAATATGTTTATTATACACGT | $P_{ugtP}$ F overlapping ori_up |
| mdb6 | **GTGAACAGCTCTTCTCCTTTTGACAT**TAATAGCCACCCTCCGTTAG | $P_{ugtP}$ R overlapping *gfp* |
| mk48 | ATGTCAAAAGGAGAAGAGCTGTTCAC | *gfp* F |
| mdb7 | **GATCCTAGGTGGGCCCAAT**TTATTTAACGAAGAATCTTGCATATAAAG | *ugtP* R overlapping *spc* |
| mk188 | ATTGGGCCCACCTAGGATC | *spc* F |
| mdb8 | AGGTGTCGACATTGGTGGTATCGCTGTTGC | ori_down R w/ SalI RS |

| | | |
|---|---|---|
| **Primers for construction of pMAD-Δ*ugtP*::*spc*** | | |
| mk501 | CAACGCCTCGCAGTCGTCC | *ugtP*_up F |
| mk502 | **TTTCCGTTAATCAAATTGCTCAT**TAATAGCCACCCTCCGTTAG | *ugtP*_up R overlapping *spc* |
| mk503 | ATGAGCAATTTGATTAACGGAAA | *spc* F |
| mk504 | CTAATTGAGAGAAGTTTCTATAG | *spc* R |
| mk505 | **CTATAGAAACTTCTCTCAATTAG**AAAATTAAGTATGCTACACAGAC | *ugtP*_down F overlapping *spc* |
| mk506 | ACGTGGATCCGATAGCTAAAGCGATAATCCAC | ugtP_down R w/ BamHI RS |
| **Primers for construction of bacterial two-hybrid (BACTH) constructs** | | |
| mk488 | GCAGGTCGAC*AGGAAACAGCT*ATGGTTACTCAAAATAAAAAGATA | pUT18-*ugtP* F w/ SalI RS and RBS |
| im228 | TAGAGGATCCTCTTTAACGAAGAATCTTGCATATAAAG | pUT18-*ugtP* R w/ BamHI RS |
| mk500 | ATCGGGATCCCGTTACTCAAAATAAAAAGATATTGA | pUT18C-*ugtP* F w/ BamHI RS |
| mdb2 | ACGTGAATTCATGATTAGCGTAATTATTTAACG | pUT18C-*ugtP* R w/ EcoRI RS |
| **Primers for construction of pRAB11-constructs** | | |
| **i**m17 | ATCGGTACC*CAATAAAACTAGGAGGAAATTTAA*ATGTTAAACAAGGTTTGGTTCC | pRAB11-*cozEa* F w/ KpnI RS and RBS |
| im18 | GATGAATTCTTAGTCCTTAACATTACTGTTTG | pRAB11-*cozEa* R w/ EcoRI RS |
| im19 | ATCGGTACC*CAATAAAACTAGGAGGAAATTTAA*ATGAATGAAAATGAAAAGAATATAAG | pRAB11-*cozEb* F w/ KpnI RS and RBS |
| im20 | GATGAATTCTTATTCAACTATTTTATTACTTTCTT | pRAB11-*cozEb* R w/ EcoRI RS |
| **Primers for construction of pMAD-*cozEa-m(sf)gfp _spc*** | | |
| mk432 | ACGTCCATGGATGTTAAACAAGGTTTGGTTCC | *cozEa* F w/ NcoI RS |
| mk433 | **GATCCTAGGTGGGCCCAAT**TTACTTATAAAGCTCATCCATGCC | *gfp* R overlapping *spc* |
| mk188 | ATTGGGCCCACCTAGGATC | *spc* F |
| mk434 | ACGTGTCGACTGGGATTAGATATTCTATCCGT | *cozEa*_down R w/ SalI RS |
| **Primers for construction of pMAD-*cozEb-m(sf)gfp _spc*** | | |
| mk435 | ACGTCCATGGATGAAAATGAAAAGAATATAAGAAAG | *cozEb* F w/ NcoI RS |
| mk433 | **GATCCTAGGTGGGCCCAAT**TTACTTATAAAGCTCATCCATGCC | *gfp* R overlapping *spc* |
| mk188 | ATTGGGCCCACCTAGGATC | *spc* F |
| mk436 | ACGTGTCGACTCGGGTGGTCTAACCATTGA | *cozEb*_down R w/ SalI RS |
| **Primers for construction of pET19b-*cozEa* and pET19b-*cozEb*** | | |
| mk508 | CGGCACTAGTCAT ATGTTAAACAAGGTTTGGTTCC | pET19b-*cozEa* F w/ SpeI RS |
| mk509 | ACCAGGATCCTTAGTCCTTAACATTACTGTTTG | pET19b-*cozEa* R w/ BamHI RS |
| mk510 | CGGCACTAGTCATATGAATGAAAATGAAAAGAATATAAGA | pET19b-*cozEb* F w/ SpeI RS |
| mk512 | ACCAGGATCCTTATTCAACTATTTTATTACTTTCTT | pET19b-*cozEb* R w/ BamHI RS |

a. The restriction sites are underlined, the overhangs are bolded, and the ribosomal binding sites are italicized.
b. F = forward primer, R = reverse primer, RS = restriction site, and RBS = ribosomal binding site.
